# Switch-Like Phosphorylation of WRN Integrates End-Resection with RAD51 Metabolism at Collapsed Replication Forks

**DOI:** 10.1101/403808

**Authors:** Valentina Palermo, Eva Malacaria, Maurizio Semproni, Benedetta Perdichizzi, Pasquale Valenzisi, Massimo Sanchez, Federica Marini, Achille Pellicioli, Annapaola Franchitto, Pietro Pichierri

## Abstract

Replication-dependent DNA double-strand breaks are harmful lesions preferentially repaired by homologous recombination; a process that requires processing of DNA ends to allow RAD51-mediated strand invasion. End-resection and subsequent repair are two intertwined processes, but the mechanism underlying their execution is still poorly appreciated. The WRN helicase is one of the crucial factors for the end-resection and is instrumental to select the proper repair pathway. Here, we reveal that ordered phosphorylation of WRN by the CDK1, ATM and ATR kinases define a complex regulatory layer that is essential for correct long-range end-resection connecting it to repair by homologous recombination. We establish that long-range end-resection requires an ATM-dependent phosphorylation of WRN at Ser1058 and that phosphorylation at Ser1141, together with dephosphorylation at the CDK1 site Ser1133, is needed for the correct metabolism of RAD51 foci and RAD51-dependent repair. Collectively, our findings suggest that regulation of WRN by multiple kinases functions as molecular switch to allow a timely execution of end-resection and repair at replication-dependent DNA double-strand breaks.

## INTRODUCTION

DNA double-strand breaks (DSBs) represent a major threat to the integrity of the genome. They can be produced by treatment with physical and chemical agents, such as DNA topoisomerases poisons camptothecin and etoposide (Nitiss, 2002), but they can be induced also at stalled replication forks after their collapse by specialized endonucleases (Dehé and Gaillard, 2017). In eukaryotes, DSBs formed during S-phase are repaired by homologous recombination (HR), however, they must be processed to yield an intermediate suitable for RAD51-mediated strand invasion (Ceccaldi et al., 2015; San Filippo et al., 2008). Processing of DSBs involves different proteins that ultimately are needed to carry out kb-long resection at the DNA ends (Mimitou and Symington, 2011; Ranjha et al., 2018; Symington and Gautier, 2011). This extensive end-resection, on one end, inhibits activation of proteins involved in the non-homologous end-joining (NHEJ) or microhomology-mediated end-joining (MMEJ) pathway of DSBs repair and, on the other hand, allows loading of HR factors (Aparicio et al., 2014; Ceccaldi et al., 2015; Chapman et al., 2012; Chiruvella et al., 2013). In addition, extensive long-range end-resection triggers activation of the ATR-mediated signalling (Flynn and Zou, 2011; Saldivar et al., 2017; Symington, 2016). Interestingly, as the end-processing at DSBs is regulated by ATM, and mostly CDK1, while repair by HR is regulated also by ATR, a switch between the two major regulatory circuits must occur (Saldivar et al., 2017; Symington, 2016). Although such a molecular switch is expected to involve assembly and disassembly of distinct molecular complexes, how exactly this switch takes place and which proteins are implicated in the process is only partially understood. Among proteins working at the interface between these two phases of DSBs repair by HR, one particularly interesting is the WRN protein (WRN). WRN is a factor involved in several processes linked to genome maintenance and its loss leads to hypersensitivity to agents inducing DSBs, in particular those acting during S-phase (Franchitto et al., 2000; Pichierri et al., 2011, 2000; Poot et al., 1999; Rossi et al., 2010). HR could be affected by loss of WRN at multiple levels as WRN activities are involved in promoting end-resection, but also in the pre-synaptic or post-synaptic metabolism of RAD51 (Franchitto and Pichierri, 2011; Rossi et al., 2010). During long-range end-resection, WRN is regulated by the CDK1 kinase and its phosphorylation is necessary for proper repair by HR (Palermo et al., 2016). However, WRN is also targeted by the ATM and ATR kinases (Ammazzalorso et al., 2010; Matsuoka et al., 2007) and is a partner of BRCA1 (Cheng et al., 2006). Notably, BRCA1 is required for accurate execution of end-resection and repair and is also targeted by the CDK1, ATM and ATR kinases (Chen et al., 2018; Symington, 2016). Hence, WRN is an ideal candidate to support a correct transition between the two phases of DSB repair by HR. Unfortunately, loss of WRN leads to complex phenotypes in response to DSBs, since its absence is offset by BLM during end-resection and HR (Nimonkar et al., 2011; Pichierri et al., 2001; Sturzenegger et al., 2014).

In this study, we took advantage from our WRN mutants to investigate on the cross-talk between CDK1- and ATM/ATR-dependent WRN regulation in response to DSB formation, and to assess if WRN regulation is involved in coordinating end-resection with initiation of HR repair. Our results demonstrate that regulation by the CDK1 and ATM kinases is an ordered process involved in establishing long-range end-resection. We show that phosphorylation by CDK1 at Ser1133 stimulates modification at Ser1058 by ATM, which is the regulatory event for correct resection by WRN. Notably, these two “pro-resection” sites must be turned off to promote ATR-dependent modification of Ser1141, which depends on long-resection, allows correct metabolism of RAD51 foci and HR. Deregulated phosphorylation by ATR leads to inappropriate modification at Ser1133 that prevents productive RAD51 foci formation and repair of DSBs.

Altogether, using regulatory mutants of WRN acting as a sort of separation-of-function forms, we unveil a critical role of WRN as a molecular switch to contribute to the passage through different stages of HR at collapse replication forks. Moreover, our findings contribute to shed further light into how post-synaptic RAD51 metabolisms is regulated in human cells in response to replication-dependent, single-ended, DSBs.

## RESULTS

### ATM-dependent phosphorylation of WRN requires prior modification by CDK1 upon replication fork collapse

To understand whether ATM could target WRN during DSBs processing, HEK293TshWRN cells were transiently transfected with wild-type Flag-WRN and the phosphorylation at S/TQ sites was evaluated by anti-pS/TQ Western blotting in anti-Flag immunoprecipitates. Our results showed that CPT treatment increased the S/TQ phosphorylation of WRN and this is more efficiently reduced by ATMi (KU-55933) than ATRi (VE-821) (Fig. 1A), suggesting that the majority of phosphorylation events depends on ATM during CPT, either directly or indirectly. Notably, phosphorylation at S/TQ motifs of WRN was substantially reduced also by the CDK inhibitor Roscovitine upon treatment (Fig. 1A). Of note, in untreated conditions, both ATR and ATM contributed similarly to the WRN modifications at ST/Q sites while the CDK inhibitor Roscovitine did not affect the level of phosphorylation at those sites (Fig. 1A). As WRN contains six S/TQ sites and two of them, Ser1141 and Ser1058, are critical for ATM-dependent regulation (Ammazzalorso et al., 2010; Matsuoka et al., 2007), we narrowed our analysis to Ser1141 for which an anti-pS1141-WRN antibody is commercially available. As shown in Figure 1B, treatment with CPT strongly enhanced S1141-WRN phosphorylation. Of note, Ser1141 phosphorylation of WRN was similarly reduced by either ATMi or ATRi and substantially decreased after Roscovitine treatment (Fig. 1B). These results suggest that CDK-dependent phosphorylation is required for the ATM/ATR-dependent regulation of WRN in response to CPT.

**Figure 1.**
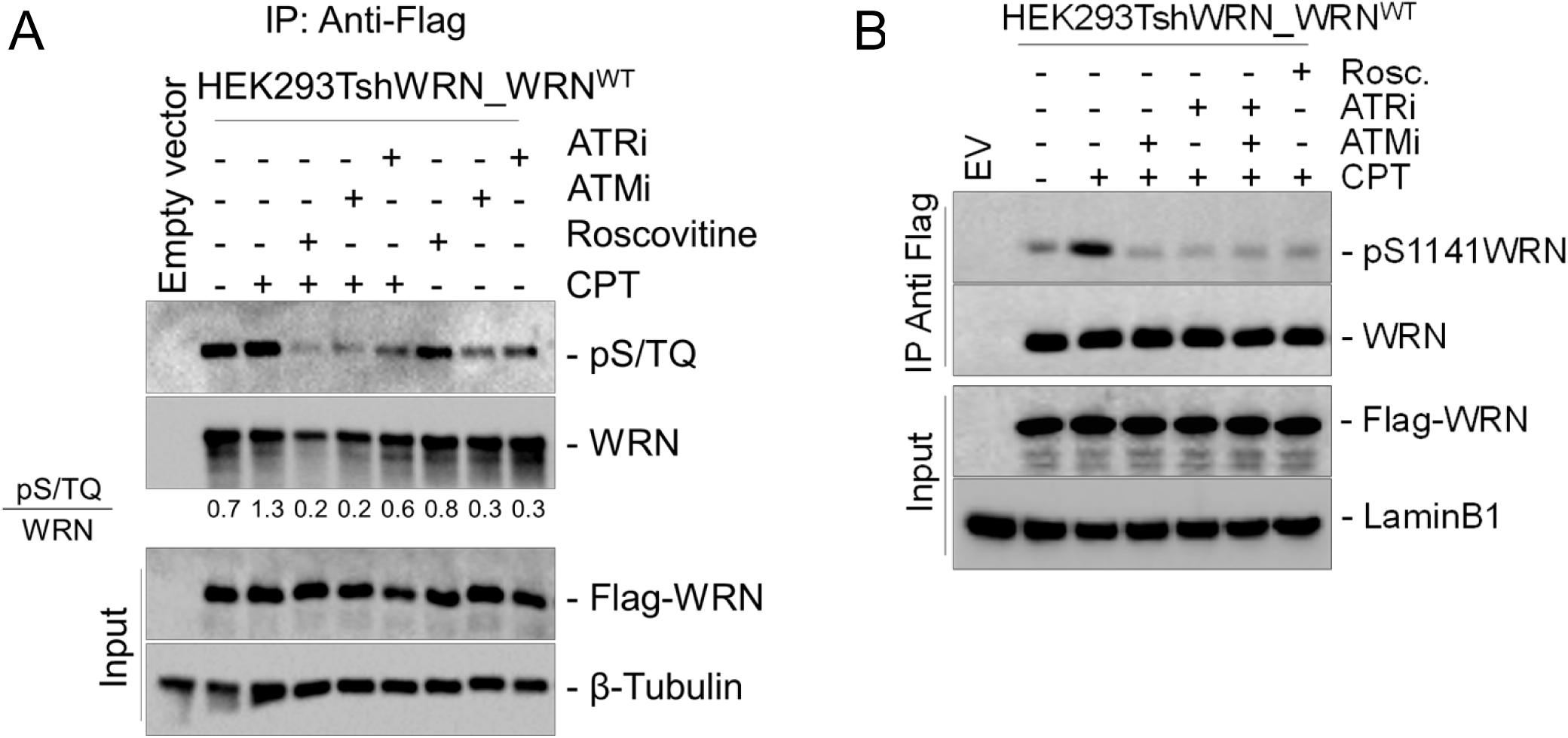
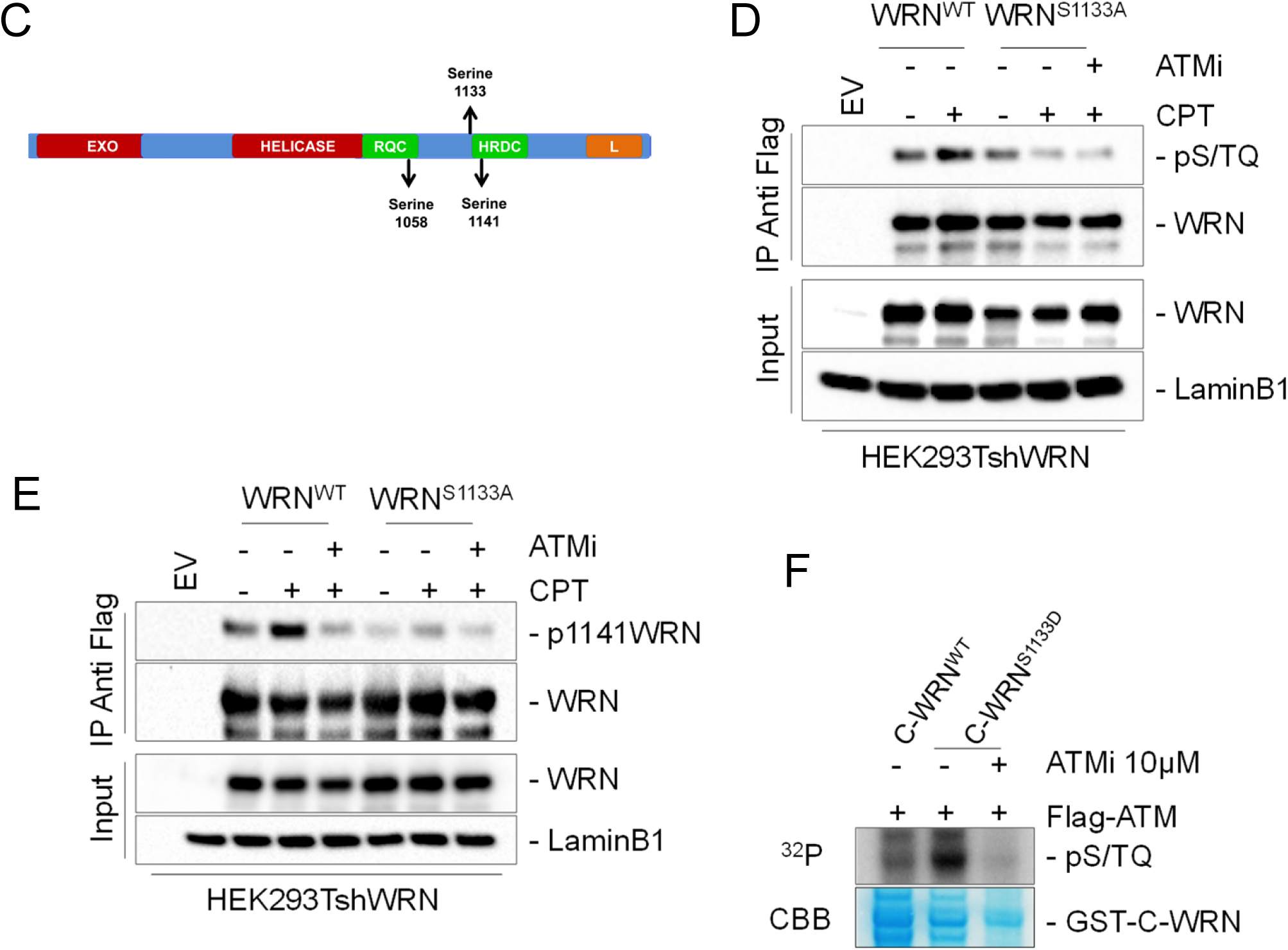
ATM/ATR-dependent WRN phosphorylation requires CDK activity upon CPT. A) WRN was immunoprecipitated from cells transiently transfected with Flag-WRN wild-type (WRN^WT^) and treated with 20μM Roscovitine, 10μM ATMi (KU-55933), 10μM ATRi (VE-821), alone or in combination, and then with 5μM CPT for 4 hours. Nine-tenth of IPs was analysed by Western Blotting (WB) with the anti-pS/TQ antibody, while 1/10 was analysed by anti-WRN antibody. Input represents 1/50 of the lysate. Anti-Flag antibody was used to verify transfection and an anti-β-Tubulin antibody was used as loading control. Quantification of the representative blots is reported below each lane. B) WRN protein picture and its different phosphorylation sites. C) Cells treated and subjected to anti-Flag-WRN IPs as in “A”. Nine-tenth of IPs was subjected to WB with an anti-pS1141WRN antibody, while 1/10 was detected by anti-Flag antibody, as indicated. Input represents 1/50 of the lysate. Anti-LaminB1 was used as loading control. D) Cells transiently transfected with WRN^WT^ or WRN^S1133A^ mutant were treated with 10μM ATMi then treated with 5μM CPT for 4 hours followed by IP/WB. Ninetenth of IPs were analysed by WB with the anti-pS/TQ antibody, while 1/10 was analysed by anti-WRN antibody, as indicated. Input represents 1/50 of the lysate. Anti-Flag was used to verify transfection and an anti-LaminB1 antibody was used as loading control. D) Cells treated and subjected to anti-Flag-WRN IPs as in “C”. Nine-tenth of IPs were analysed by WB with the anti-pS1141WRN antibody. F) In vitro ATM kinase assay. For kinase assay, 2 μg of immunopurified GST-tagged WRN wild-type fragment (C-WRN^WT^) or WRN phosphomimetic mutant fragment (C-WRN^S1133D^) were phosphorylated in vitro using Flag-tagged ATM kinase, immunoprecipitated with specific anti-Flag-conjugated beads. Immunoblotting was used to analyse ATM-dependent phosphorylation level in different WRN fragments using an anti-pS/TQ antibody. Treatment with 10μM ATM inhibitor (KU-55933) was used as a control, as indicated. Coomassie (CBB) showed GST-C-WRN fragments in the gel, as a control.

Ser1058 and Ser1141 are close to the regulatory CDK site Ser1133 (Fig. 1C). To determine if reduction of ATM/ATR-dependent phosphorylation of WRN upon CDK inhibition could depend specifically on loss of Ser1133 phosphorylation, we examined WRN phosphorylation at S/TQ sites by IP/WB experiments in cells expressing the CDK1-unphosphorylable S1133A mutant (Palermo et al., 2016). We found that S/TQ phosphorylation was largely suppressed in the S1133A-WRN mutant in response to CPT but not under unperturbed conditions when compared with cells expressing WRN wild-type (Fig. 1D). Consistently, the S1133A-WRN mutant also showed a strong reduction of phosphorylation at Ser1141 under all conditions (Fig. 1E)

As the majority of phosphorylation events on WRN S/TQ sites depends on ATM during CPT and that S/TQ phosphorylation is largely suppressed in the S1133A-WRN mutant, we investigated if phosphorylation of S1133 improved the ability of ATM to modify WRN. Thus, we performed an ATM *in vitro* kinase assay using a C-terminal fragment of WRN as substrate, either wild-type or containing the S1133D phosphomimetic mutation. As shown in Figure 1F, the S1133D phosphomimetic mutation greatly increased phosphorylation by ATM. These results suggest that CDK1 primes ATM-dependent modification of WRN at its target sites.

Of note, the CDK1-dependent phosphorylation of WRN at S1133 was dependent on ATM activity (Suppl. Fig. 1A), and ATM/ATR- and CDK1-dependent phosphorylation of WRN were found to be both MRE11 nuclease-dependent (Suppl. Fig. 1B).

Altogether, our results indicate that the response to single-ended DSBs (seDSBs), as induced by CPT treatment, requires an ordered phosphorylation of WRN by CDK1 and ATM/ATR. They also show that ATM and MRE11-dependent events are upstream to the CDK1-dependent phosphorylation of WRN, which, in turn, is required for ATM/ATR-dependent modification of the protein.

### Regulated and site-specific phosphorylation of WRN is crucial for end-resection at DSBs

To determine whether ATM-dependent phosphorylation of WRN could be involved in the resection of DNA ends, we evaluated formation of ssDNA by the IdU/ssDNA assay (Fig. 2A). End-resection was assessed in WS cells stably expressing the wild-type WRN (WRN^WT^), its ATM-unphosphorylable (WRN^3AATM^) or phosphomimetic (WRN^3DATM^) mutant and, as a control, the end-resection-defective WRN mutant (WRN^S1133A^) (See Supplementary Fig. 2 for expression levels). Analysis was performed at 90mins of treatment with CPT, when formation of ssDNA is dependent on S1133 phosphorylation of WRN (Palermo et al., 2016). Loss of CDK1-dependent phosphorylation of WRN reduced formation of ssDNA upon CPT exposure, as expected (Fig. 2B). Mutations abrogating ATM-dependent phosphorylation of WRN significantly decreased ssDNA exposure, which was surprisingly lowered also by mutations mimicking constitutive phosphorylation by ATM on WRN (Fig. 2B). Inhibition of ATM completely suppressed ssDNA in all cell lines, consistent with ATM targeting multiple proteins during end-resection (Supplementary Fig. 3A).

**Figure 2.**
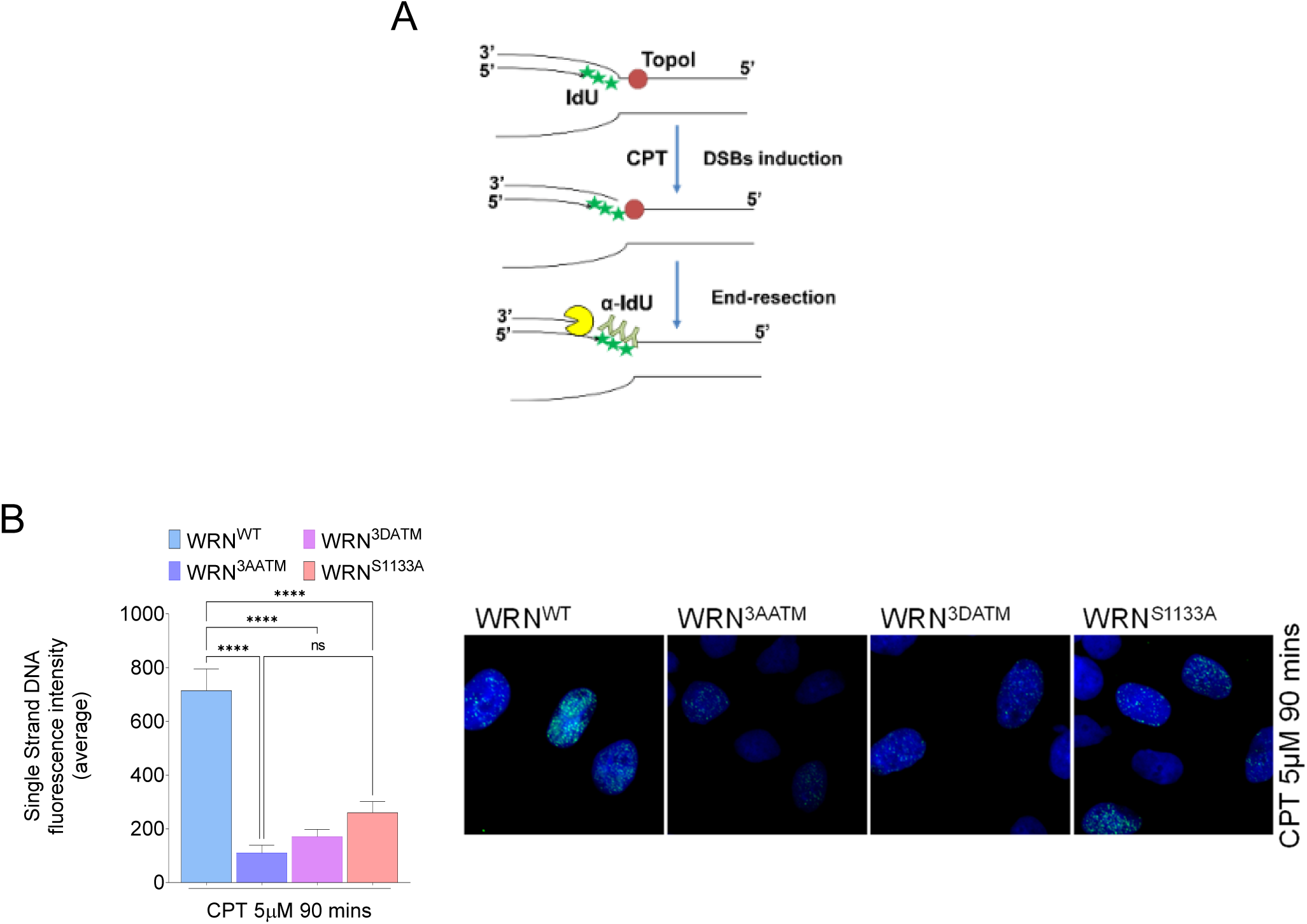
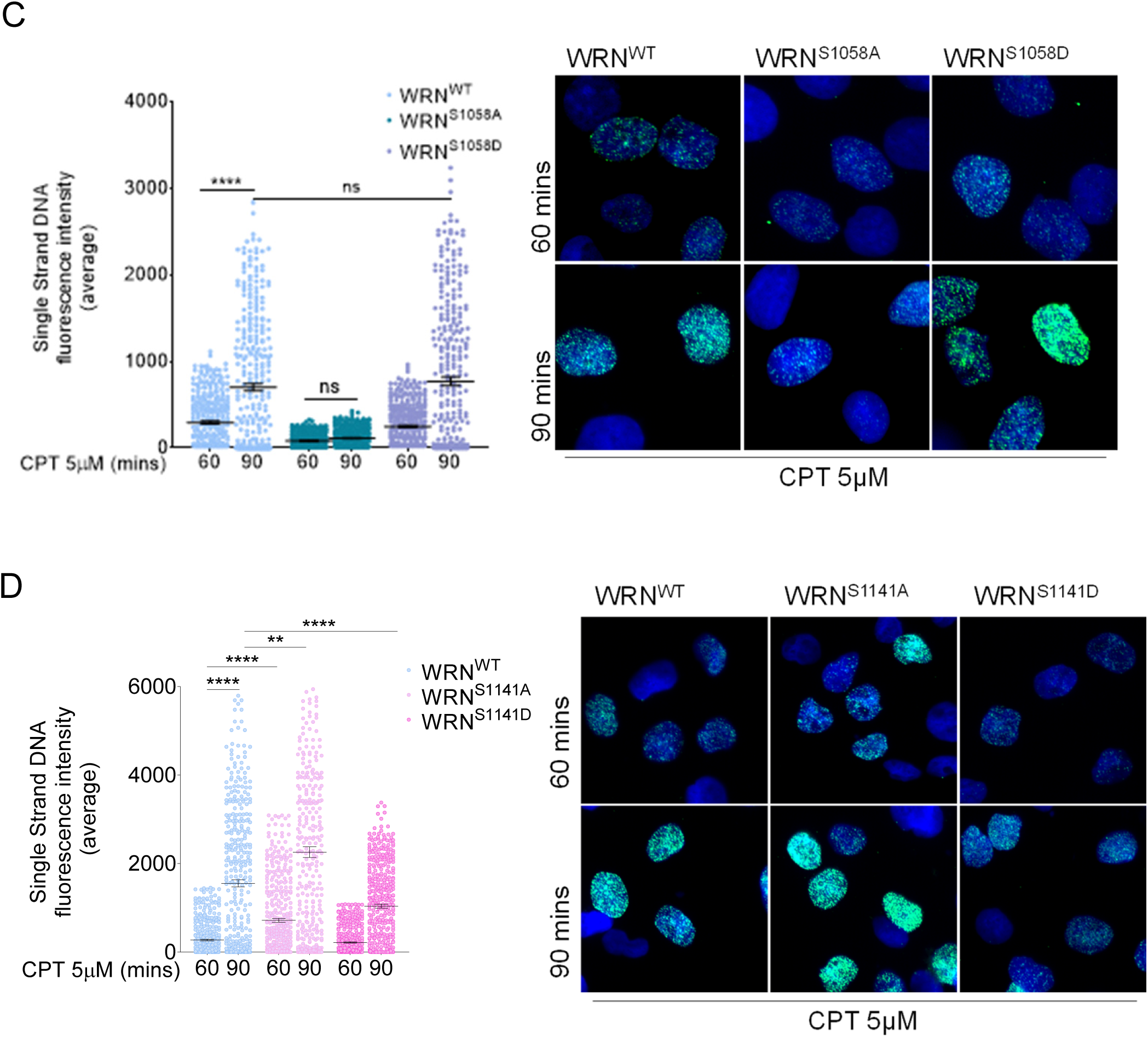
Phosphorylation by ATM/ATR of WRN at distinct sites differently affects end-resection of DSBs. A) Cartoon depicting the ssDNA assay by native anti-IdU immunofluorescence. The scheme shows how ssDNA can be visualized at collapsed replication forks. CPT treatment results in one-ended DSBs at replication forks leading to 5′-3′ resection of template DNA by nucleases (pacman) thus exposing nascent ssDNA, which is detected by our native IdU/ssDNA assay. Nascent DNA was pre-labelled for 15 min with IdU before treatment and labelling remained during treatment with CPT. B) WS-derived cell lines complemented with different WRN mutants were labelled, treated with CPT and IdU/ssDNA assay was performed. WB shows WRN expression in different cell lines using anti-WRN antibody. The graph shows the mean intensity of IdU/ssDNA staining for single nuclei measured from three independent experiments (*n*=300, each biological replicate), data are presented as mean±SE. Representative images of IdU/ssDNA-stained cells are shown. Statistical analysis was performed by the ANOVA test (**** = p<0.0001; ** = *P*<0.01; ns = not significant). C-D). WS fibroblasts were transiently transfected with the indicated WRN-expressing plasmid. WB shows WRN expression levels 48hrs after transfection using anti-WRN antibody. The ssDNA was analysed at different time points, as indicated. The dot plots show the mean intensity of IdU/ssDNA staining for single nuclei (n=300, two biological replicates). Data are presented as mean±SE. Representative images of IdU/ssDNA-stained from CPT-treated cells are shown. Statistical analysis was performed by the ANOVA test (**** = p<0.0001; ** = *P*<0.01; * = *P*>0.05; ns = not significant).

The similar phenotype shown by cells expressing the ATM unphosphorylable or phosphomimetic triple WRN mutant prompted us to explore the functional relevance of each single phosphorylation site. Single unphosphorylable (S>A) or phosphomimetic (S>D) WRN mutants were transiently expressed in WS cells and the ability to perform end-resection was evaluated by the ssDNA/IdU assay at 60 and 90mins of CPT treatment. While mutation of the S1292 site was functionally irrelevant for the end-resection (Suppl. Fig. 3B), expression of the S1058A-WRN mutant significantly impaired end-resection and recapitulated the phenotype of WRN^3AATM^ cells (Fig. 2B, C). Compared to the wild-type, expression of the phosphomimetic S1058D-WRN mutant did not affect formation of ssDNA (Fig. 2C), thus diverging from the end-resection defect observed in WRN^3DATM^ cells.

In sharp contrast with the S1058A-WRN mutant, the S1141A mutant was end-resection proficient and, at the early time-point after CPT treatment, showed also a slightly enhanced level of ssDNA compared to the wild-type (Fig. 2D). Of note, the defective end-resection phenotype of the S1058A-WRN mutant was detectable also after treatment with the radiomimetic drug Bleomycin, while the phenotype of S1141A-WRN was not detectable after this treatment (Supplementary Fig. 4A). In contrast, all the mutants exhibiting altered nascent ssDNA exposure, as a redout of end-resection, in response to replication-dependent DSBs did not induce any alteration in the formation of ssDNA in HU-treated cells (Supplementary Fig. 4B). Collectively, these results suggest that phosphorylation at these sites is not involved in regulating degradation of intact forks but are important to process DSBs.

Worth noting, however, also cells expressing the phosphomimetic S1141D-WRN mutant apparently showed a reduced end-resection in CPT. To further investigate the phenotype observed in the S1141-WRN mutant, we performed quantitative evaluation of end-resection at single DSB sites introduced by Cas9/sgRNA (Dibitetto et al., 2018). To this end, we transiently transfected WS cells with plasmid expressing the wild-type WRN or its S1141A mutant and plasmids expressing Flag-Cas9 and sgRNA guides against one AsiSI site (Supplementary. Fig. 5). Formation of ssDNA at Cas9-induced DSB was analysed by droplet-digital PCR (ddPCR) at increasing distance from the cutting site and normalized against the cut efficiency (Dibitetto et al., 2018). As shown in Figure 3A, although loss of Ser1141 phosphorylation of WRN increased the number of resected ends at each of the distances analysed, the effect was most evident at the most distant site. These data, together with the ssDNA/IdU assay, suggest that phosphorylation of WRN at Ser1141 limits somehow the processing of DNA ends. Unexpectedly, considering the reduced detection of ssDNA in the ssDNA/IdU assay (Fig. 2D), the phosphomimetic S1141D WRN mutant showed increased levels of ssDNA in the Cas9/sgRNA resection assay (Fig. 3A).

**Figure 3.**
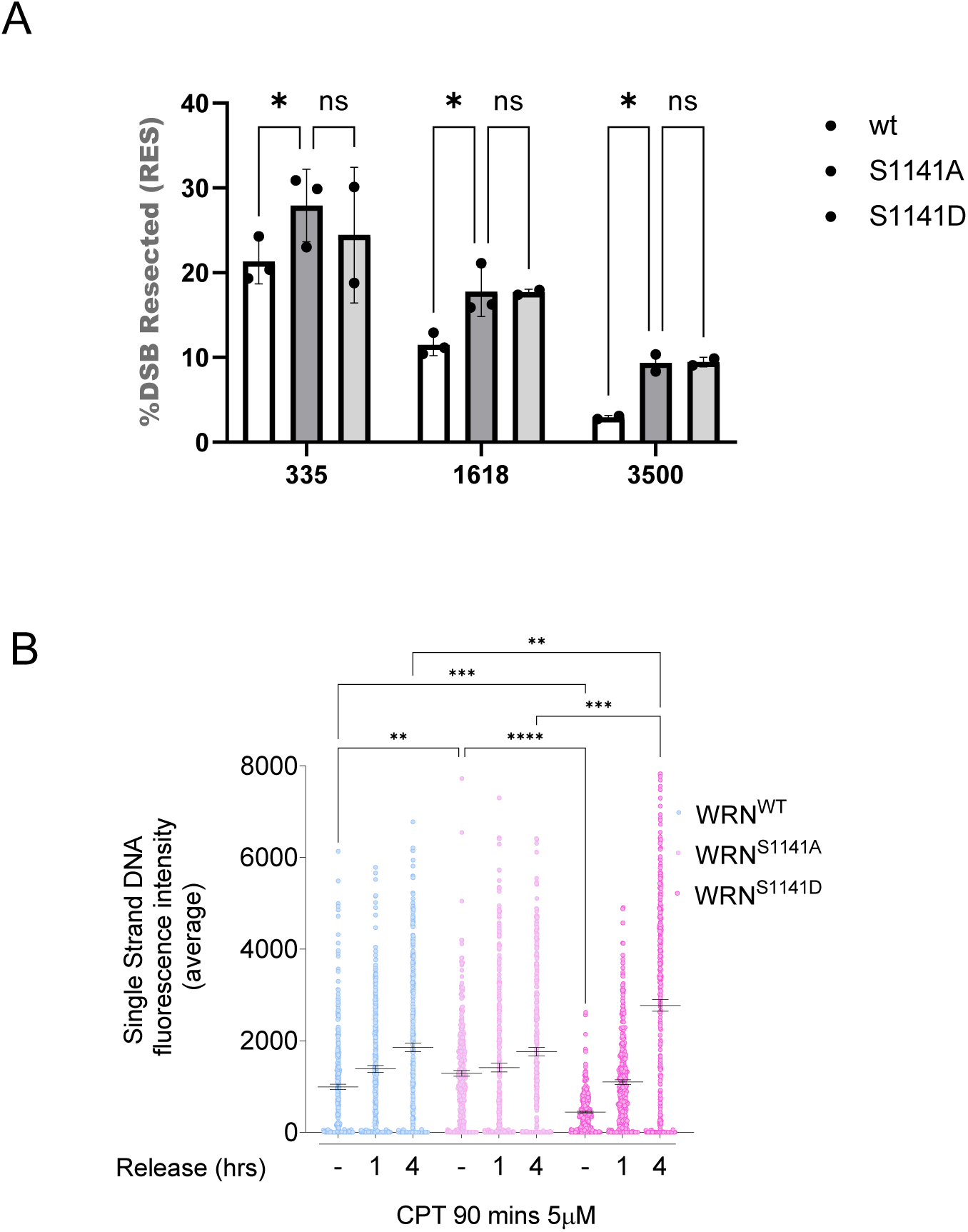
Altered phosphorylation of WRN at S1141 affects end-resection. A) Graph shows the analysis and the quantification of Cas9-induced and processed DSBs in WS fibroblast transiently transfected with WRN mutants, as indicated. Data are represented as the percentage of DSBs resected on a specific chromosomal site. B) WS-derived cell lines complemented with different WRN mutants were labelled and treated with CPT followed by different time points of release in free medium before performed IdU/ssDNA assay. Dot plot shows the mean intensity of IdU/ssDNA staining for single nuclei (n=300, two biological replicates). Data are presented as mean±SE. Statistical analysis was performed by the ANOVA test (**** = p<0.0001; ** = *P*<0.01; * = *P*>0.01; ns = not significant).

Since the ssDNA/IdU assay was performed very early during CPT treatment, while the Cas9/sgRNA assay was presumably performed at a later stage, we reasoned that the unexpected phenotype of S1141D-WRN mutant in the Cas9/sgRNA might be related to aberrant processing at an advanced stage of DSBs repair. Thus, we evaluated formation of ssDNA at the nascent strand during recovery from CPT treatment. Formation of ssDNA was analysed by the IdU/ssDNA assay at 0, 1 and 4h of recovery from a 90mins treatment with CPT (Fig. 3B). In wild-type cells, formation of nascent ssDNA was detected also during recovery and increased with time. Cells expressing the S1141A-WRN mutant showed enhanced formation of ssDNA compared to the wild-type during CPT, although the level of ssDNA remained stable with time during recovery. Surprisingly, while slightly reduced levels of ssDNA were detected in the S1141D-WRN mutant during treatment, a striking increase was observed during recovery, when levels of ssDNA significantly exceeded those detected in the wild-type recapitulating what observed in the Cas9/sgRNA assay. In both wild-type and S1141D WRN formation of nascent ssDNA was dependent on DNA2, suggesting that it represents aberrant long-range resection (Supplementary Fig. 6).

Collectively, these findings suggest that phosphorylation at Ser1058 by ATM, together with that performed by CDK1 at Ser1133, is critical for end-resection, while phosphorylation status of Ser1141 is important for its regulation.

### Ser1141 and Ser1133 of WRN are inversely regulated during distinct stages of HR

We found that regulated phosphorylation of WRN at Ser1141 may be involved in controlling the resection of DSBs during the repair stage of HR. Thus, we tested the possibility that modification of Ser1141 and Ser1133 could be differently regulated during separate stages of HR. To test this hypothesis, we performed IP/WB to evaluate Ser1133 or Ser1141 phosphorylation at 1h and 4h of treatment (Fig. 4A), which corresponds to end-resection stage and initial recruitment of RAD51, or after recovery, when DSBs are repaired (Supplementary Fig. 7A and B;(Palermo et al., 2016)). Consistent with its role during end-resection, phosphorylation at Ser1133 was detectable already at 1h of treatment and remained elevated at 4h (Fig. 4A). However, phosphorylation at Ser1133 of WRN declined after recovery (Fig. 4A), when formation of RAD51 foci increased strongly (Supplementary Fig. 6B). In contrast, phosphorylation at Ser1141 was barely detectable at 1h of treatment, increased strongly at 4h and remained elevated also during recovery from CPT (Fig. 4A). Engagement of the post-resection stage of HR has been correlated with ATR-dependent phosphorylation (Buisson et al., 2017). During 24h of recovery from CPT, Ser1141 can be phosphorylated by ATR (Su et al., 2015). Thus, we wondered whether ATR could target Ser1141 also soon after recovery. Thus, we analysed the level of Ser1141 phosphorylation in cells exposed to CPT and treated with the ATRi, the ATMi or both during a 4h recovery (Fig.4B) to maximise differences observed at 2h (Fig. 4A). At the same time, we evaluated the level of Ser1133 phosphorylation, a readout of the pro-resection role of WRN (Palermo et al., 2016).As expected, phosphorylation of Ser1141 increased during recovery, while phosphorylation of Ser1133 was elevated during treatment but decreased thereafter (Fig. 4B). Surprisingly, during recovery, inhibition of ATM only barely affected phosphorylation of Ser1141, while inhibition of ATR reduced its level drastically (Fig 4B).

**Figure 4.**
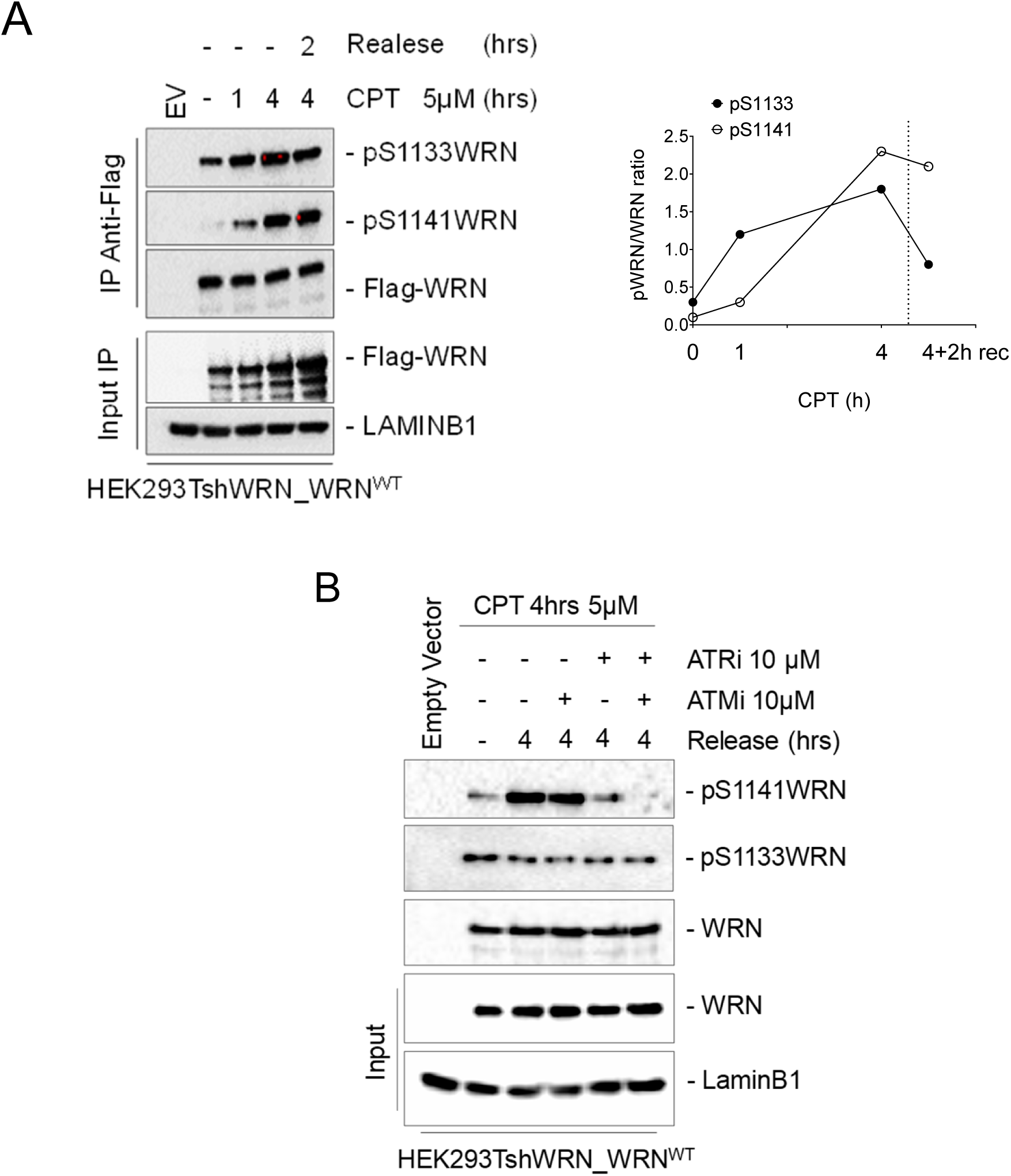
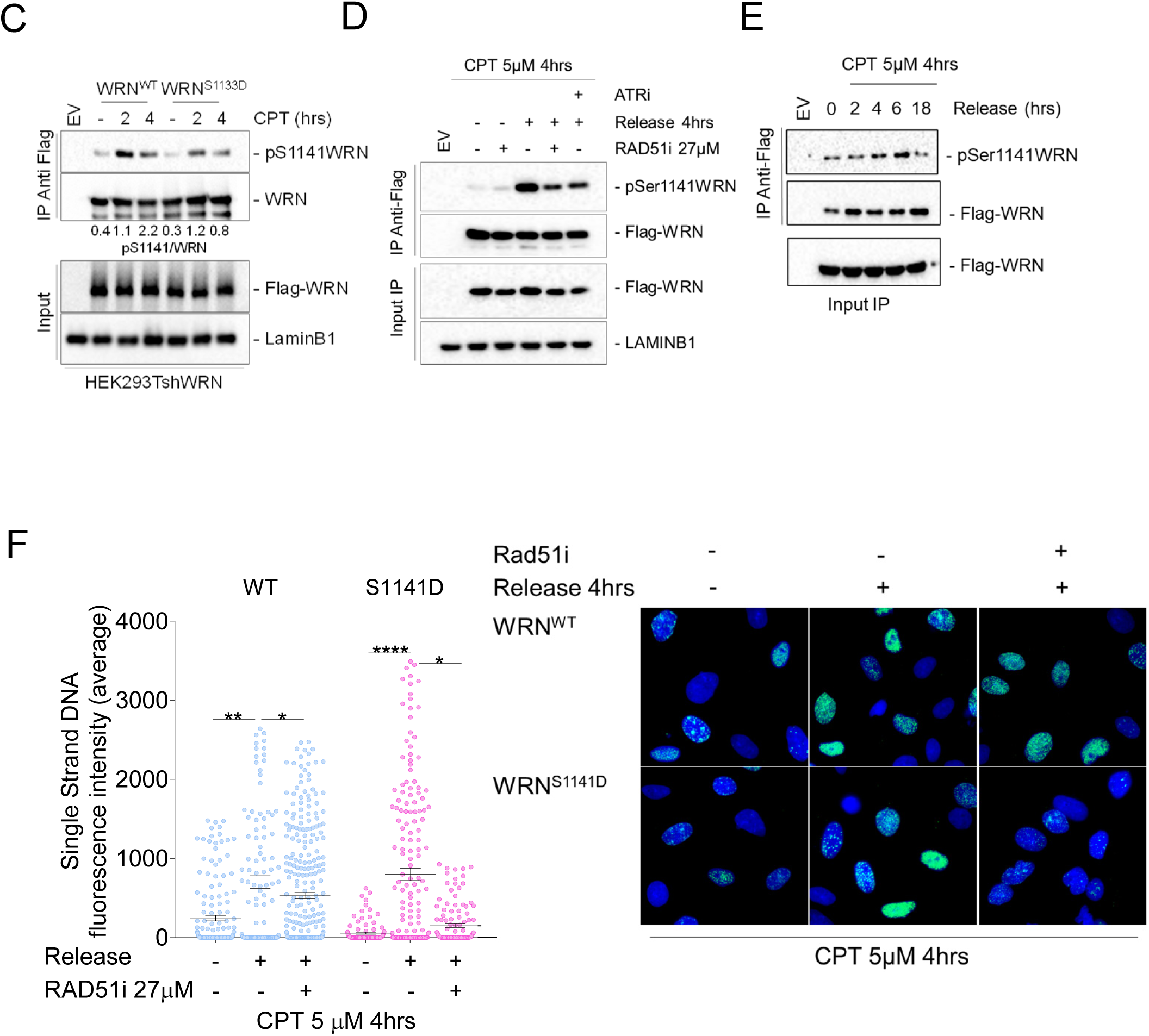
Phosphorylation of WRN by ATM/ATR occurs at the end of resection and requires its correct execution. A) Cells were transiently transfected with an empty vector or a vector expressing Flag-tagged WRN wild-type (WRN^WT^) and were treated with 5μM CPT for different time points as indicated. In addition, treated cells were recovered for 2 hours in drug-free medium. Flag-WRN was immunoprecipitated and 9/10 of IPs was analysed by WB with both the anti-pS1133WRN and the anti-pS1141WRN antibodies, while 1/10 was detected by anti-WRN, as indicated. One-fiftieth of the lysate was blotted with an anti-Flag antibody to verify transfection. An anti-LaminB1 antibody was used as loading control. The graph show kinetics of WRN phosphorylations levels and data are from the representative experiment. B) Cells were transiently transfected as in “A”. Cells were treated with 5μM CPT for 4hrs. Cells were recovered in CPT-free medium and treated with 10μM ATMi (KU-55933), 10μM ATRi (VE-821), alone or in combination. Flag-WRN was immunoprecipitated and IPs was analysed and blotted as indicated in “A”. C) Cells were transiently transfected with an empty vector or a vector expressing Flag-tagged WRN wild-type (WRN^WT^) or its phosphomimetic mutants (WRN^S1133D^) and treated with 5μM CPT for different time points as indicated. Flag-WRN was immunoprecipitated and 9/10 of IPs was analysed by WB with both the anti-pS1141WRN antibody, while 1/10 was detected by anti-WRN, as indicated. One-fiftieth of the lysate was blotted with an anti-Flag antibody to verify transfection. An anti-LaminB1 antibody was used as loading control. D) Cells were transiently transfected as in “A”. Cells were treated with 5μM CPT for 4hrs in combination or not with 27 μM RAD51 or 10 μM ATRi inhibitor and were recovered for 4 hours in CPT-free medium. Flag-WRN was immunoprecipitated and IPs was analysed and blotted as indicated in “C”. E) Cells were transiently transfected as in “A”, treated with 5μM CPT for 4hrs and were recovered for different time points, as indicated. Flag-WRN was immunoprecipitated and IPs was analysed and blotted as indicated in “C”. F) WS-derived cell lines complemented with different WRN mutants were labelled, treated with CPT and IdU/ssDNA assay was performed. The graph shows the mean intensity of IdU/ssDNA staining for single nuclei measured from three independent experiments (*n*=300, each biological replicate), data are presented as mean±SE. Representative images of IdU/ssDNA-stained cells are shown. Statistical analysis was performed by the ANOVA test (**** = p<0.0001; ** = *P*<0.01; * = *P*>0.05).

To further demonstrate that phosphorylation of Ser1133 by CDK1 and that of Ser1141 by ATR are inversely regulated, we analysed phosphorylation at Ser1141 after 2 or 4h of CPT treatment in cells transfected with the wild-type or the S1133D WRN mutant. As shown in Figure 4C, the presence of the phosphomimetic S1133D mutation halved the amount of Ser1141 phosphorylation after 4h of CPT. Moreover, phosphorylation by ATR at Ser1141 was impaired in cells expressing the unphosphorylable S1133A WRN mutant or treated with the ATMi (Supplementary. Fig. 8), which both impair long-range end-resection (Palermo et al., 2016). Consistent with failure to initiate end-resection, both conditions strongly reduced modification of WRN at Ser1141 also during recovery (Supplementary Fig. 8). This result shows that ATR targets Ser1141 of WRN even early after recovery and suggests that the two phosphorylation events represent molecular switches for two different roles of WRN during the resection or the repair stages of seDSBs by HR. Moreover, taken together, these data indicate that ATR targets Ser1141 of WRN after initiation of long-range resection and mostly when it is completed.

To determine whether phosphorylation of WRN at S1141 could correlate with the initiation of repair phase of seDSBs by HR, that is after initiation of strand-invasion, we blocked formation of RAD51 nucleofilaments with the B02 inhibitor (RAD51i) and analysed the level of WRN phosphorylation by IP/WB in CPT and during recovery (Fig. 4D). As expected, Ser1141 phosphorylation of WRN was lower during treatment and was not affected by inhibition of RAD51. However, inhibition of RAD51 nucleation greatly reduced Ser1141 phosphorylation during recovery from CPT, suggesting that this modification of WRN depends on strand invasion. To assess if phosphorylation of Ser1141 of WRN was turned off with time during recovery, we performed IP/WB assays at multiple time points after release from CPT. As shown in Figure 4E, Ser1141 phosphorylation increased at 4 and 6h of recovery but drastically decreased at 18h of recovery when most, if not all, DSBs are repaired (Palermo et al., 2016).

Thus, we reasoned that aberrant exposure of nascent ssDNA observed in the S1141D-WRN mutant could be correlated with the metabolism of the RAD51 filaments during RAD51-dependent repair. If this hypothesis was true, aberrant exposure of nascent ssDNA should be reduced by inhibition of strand invasion consistent with the relationship with RAD51 nucleation of the S1141 modification (Fig. 4D). To verify this hypothesis, we analysed nascent ssDNA in cells treated with the RAD51i to block RAD51 nucleofilament formation during recovery from CPT (Fig. 4F). RAD51 inhibition slightly reduced formation of nascent ssDNA in wild-type cells, however, it suppressed the strong increase of ssDNA in the S1141D-WRN mutant.

Collectively, these data indicate that phosphorylation at Ser1133 and Ser1141 of WRN are inversely regulated and suggest that modification of Ser1141 by ATR occurs immediately after long-range end-resection, increases during strand-invasion, and then needs to be turned off to avoid excessive accumulation of RAD51-dependent ssDNA.

### Correct metabolism of RAD51 foci and repair by HR require regulated phosphorylation of WRN at Ser1141

Our data indicates that phosphorylation of WRN by ATR at Ser1141 occurs during formation of RAD51 nucleofilaments. These findings led us to analyse whether phosphorylation at Ser1141 could affect the dynamics of RAD51 foci after recovery from CPT (Fig. 5A). No significant differences were observed in the ability to form RAD51 foci during CPT between cells expressing WRN wild-type or mutated at S1141 (Fig. 5A). During recovery, in wild-type cells, formation of RAD51 foci increased at 1h and decreasing thereafter. Inability to phosphorylate Ser1141 of WRN apparently altered the kinetics of RAD51 foci formation, which was apparently delayed, in agreement with more end-processing. By contrast, the number of RAD51-positive cells and the brightness of foci were greatly increased and sustained during recovery in the S1141D WRN mutant. At collapsed forks, RAD51 is loaded at the ssDNA exposed in the nascent strand but RAD51 can be recruited also at parental ssDNA at blocked forks. To further confirm that increased recruitment of RAD51 occurred at collapsed forks, we evaluated the presence of RAD51 at nascent ssDNA by PLA (Iannascoli et al., 2015). As expected, RAD51 was found associated with nascent ssDNA in wild-type cells during recovery from CPT treatment and, consistently with IF analysis, this association was increased in the S1141D-WRN mutant (Fig. 5B and C).

**Figure 5.**
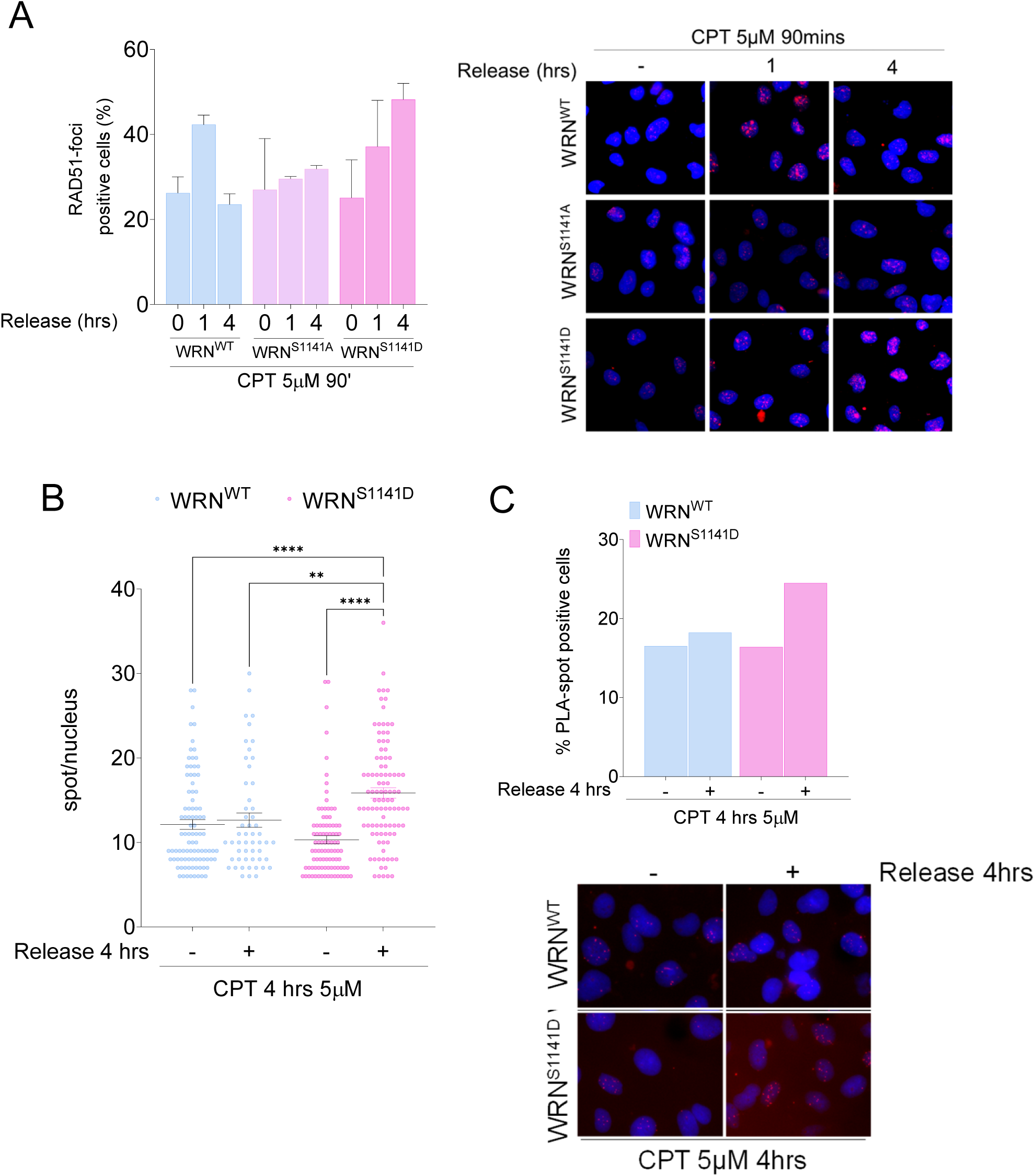
Regulated phosphorylation of WRN at S1141 is involved in RAD51 localisation. A) Cells stably expressed Flag-tagged WRN mutants were treated with 5μM CPT and allowed to recover for different time points as indicated. Then RAD51-foci staining was analysed by IF. The graph shows the percentage of RAD51-foci positive cells measured from two independent experiments (n=200, each biological replicate).data are presented as mea±SE. Representative images of RAD51 staining in response to treatment are shown. B) Analysis of ssDNA/RAD51 interaction by in situ PLA. Cells were treated with 5μM CPT and allowed to recover for 4hrs, then subjected to PLA using anti-IdU and anti-RAD51 antibodies. The panel shows representative PLA images showing association of ssDNA with RAD51, scale bar 10µm. The dot plot shows the PLA spots per PLA positive cells. At least 100 nuclei were analysed for each experimental point from two independent experiments. Values are presented as mean±SE. Statistical analysis was performed by the ANOVA test (**** = p<0.0001; ** = *P*<0.01, each biological replicate). C) The graph shows the number of PLA positive cells. At least 100 nuclei were analysed for each experimental point. Representative images from the neutral Comet assay are shown.

Consistent with aberrant and RAD51-dependent accumulation of nascent ssDNA when WRN cannot be dephosphorylated at S1141, we next investigated whether high levels of RAD51 foci represented unproductive recombination events leading to defective HR. Thus, we examined the ability to perform RAD51-dependent repair by gene conversion using a reporter assays (Fig. 6A) (Palermo et al., 2016; Pierce et al., 2001). The pDRGFP HR reporter was transiently transfected in HEK293TshWRN cells together with a plasmid expressing the wild-type, S1141A or S1141D variants of WRN and a plasmid expressing the I-SceI endonuclease. As an internal control, we also analysed HR efficiency in cells expressing the S1133A-WRN mutant, which is known to be resection and HR-defective (Palermo et al., 2016). At 72 h post-transfection, repair was evaluated by flow cytometry analysing the number of GFP-positive cells. As shown in Figure 6A, the ability to repair by HR is comparable to the wild-type in cells expressing the S1141A-WRN mutant, however it is compromised in cells expressing the S1141D-WRN or the S1133A mutants.

**Figure 6.**
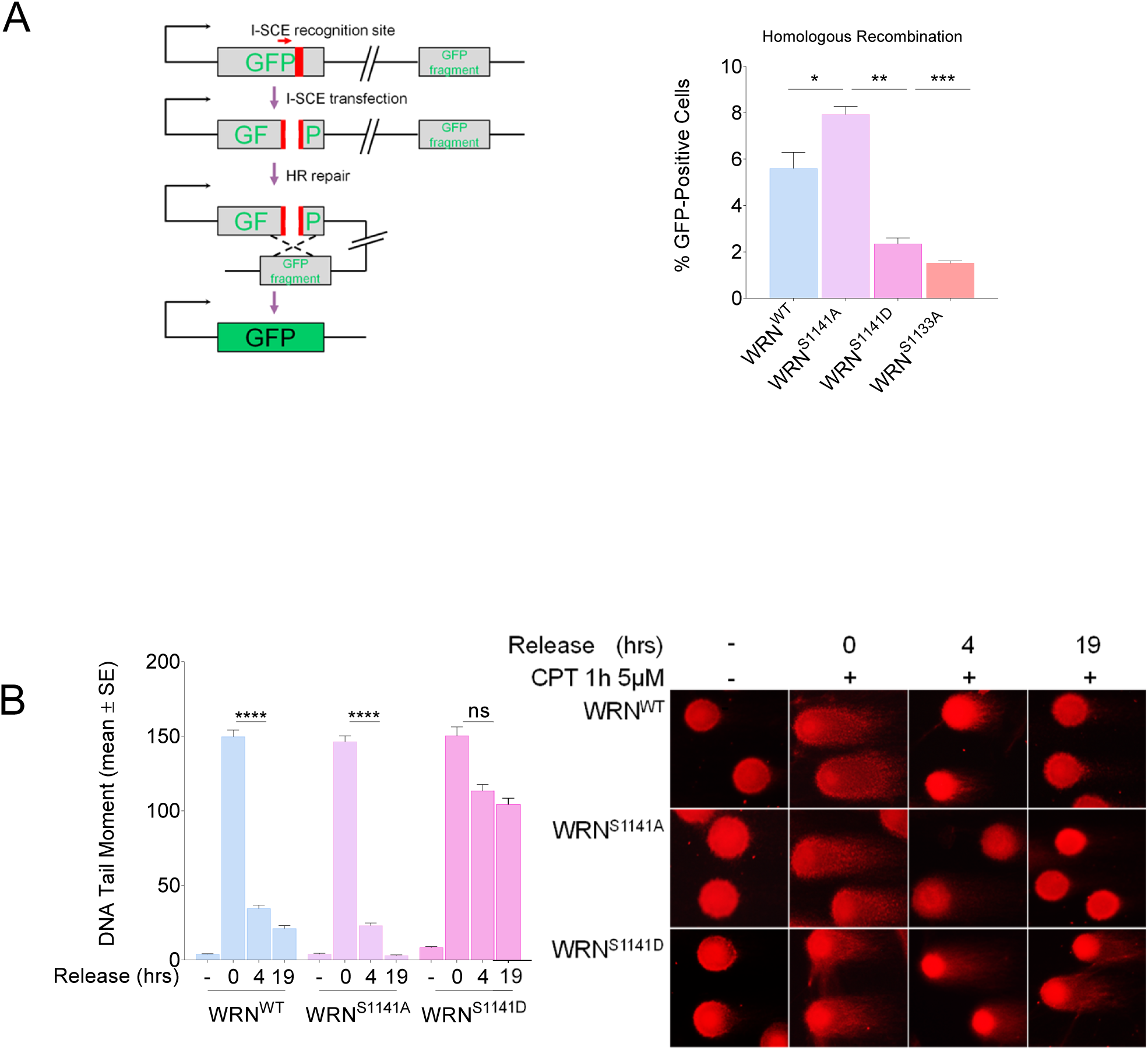
Regulated phosphorylation of WRN at S1141 is essential for DSBs repair. A) Analysis of efficiency of HR-mediated repair by reporter assay. HEK293TshWRN cells were co-transfected with the indicated WRN forms, the I-SceI expression vector pCBASce and the pDRGFP HR reporter plasmid, as described in Methods. The graph shows the percentage of GFP positive cells measured by flow cytometry. Data are presented as mean±SE from three independent experiments (ns = not significant; * = *P*<0.05; ** = *P*<0.01; *** = *P*<0.001; ANOVA test; *n*=3 × 10^5^ events each biological repeat). B) DSBs repair efficiency analysis. WS-derived SV40-trasformed fibroblasts stably expressed Flag-tagged WRN mutants were treated with 5 μM CPT for 1 h and then release in drug free medium at different time points. DSBs repair was evaluated by the neutral Comet assay. In the graph, data are presented as mean tail moment±SE from three independent experiments. Representative images from the neutral Comet assay are shown. Statistical analysis was performed by the ANOVA test (**** = p<0.0001; ns = not significant, each biological replicate).

Then, we wanted to determine if deregulated phosphorylation of WRN S1141 could affect repair of CPT-induced DSBs. Indeed, abrogation of Ser1133 phosphorylation of WRN impairs HR but induces a switch of pathway triggering repair by end-joining (Palermo et al., 2016). To this aim, we explored DSBs repair by neutral Comet assay during recovery from a short CPT treatment (Fig. 6B). As expected, in WRN^WT^ cells, DSBs were repaired almost completely at 4h of recovery and abrogation of S1141 phosphorylation (WRN^S1141A^) did not have any significant effect except a more efficient repair at 19h of recovery. Confirming the predominant engagement of HR over other repair pathways, repair was greatly reduced by the RAD51i but not by DNA-PK or LigI/III inhibition, which blocks, respectively, NHEJ or alt-EJ (Supplementary Fig. 9A, B). Conversely, WRN^S1141D^ cells showed a compromised DNA repair with only a minor fraction of DSBs being repaired even at 19h of recovery. This residual fraction of repair was completely dependent on RAD51 (Supplementary Fig 9C) and repair was rescued by abrogation of the end-resection through depletion of BRCA1 (Supplementary Fig. 9D), which redirects DSBs repair from HR to NHEJ.

Therefore, these findings indicate that switching off the ATR-dependent phosphorylation of WRN at Ser1141 is required to promote a correct post-synaptic metabolism of RAD51 foci, preventing aberrant accumulation of RAD51 foci and ensuring repair by HR.

### A timely switch from CDK1- to ATR-dependent phosphorylation of WRN is required for correct DSBs metabolism and repair by HR

Our results suggest that ATR-dependent phosphorylation of Ser1141 of WRN must take place at a specific window of opportunity to promote proper RAD51 metabolism and repair by HR. Persistent or untimely modification of Ser1141, as mimicked by the phosphomimetic mutant S1141D, leads to aberrant exposure of ssDNA and defective repair of seDSBs. Since CDK1 and ATR-dependent phosphorylation of WRN are ordered events, we hypothesized that the presence of the phosphomimetic mutation at Ser1141 might promote untimely phosphorylation at the adjacent Ser1133 in a moment in this modification should be switched off because of the low CDK1 activity. Hence, we first tested whether the presence of phosphorylated Ser1141 could affect Ser1133 phosphorylation *in vitro*. A small fragment of WRN containing the wild-type Ser1141 residue or the S>D phosphomimetic mutation was used as substrate for CDK2/Cyclin complex and phosphorylation at Ser1133 was analysed by Western blotting (Fig. 7A). Consistently with our previous data (Palermo et al., 2016), the CDK2/Cyclin complex efficiently phosphorylated Ser1133 of WRN. However, CDK/Cyclin-mediated phosphorylation of Ser1133 was greatly stimulated by the S1141D mutation. This suggested that constitutive phosphorylation of Ser1141 disrupts the ordered modifications of WRN in response to DSBs, promoting modification of Ser1133 even when CDK1 activity levels are low. Thus, we monitored the phosphorylation of Ser1133 by IP/WB in cells expressing the wild-type or the S1141D-WRN protein during CPT treatment or recovery (Fig. 7B). As expected, Ser1133 phosphorylation prevails during treatment but is extremely reduced at 2h of recovery. By contrast, mimicking a constitutive phosphorylation by ATR led to sustained Ser1133 phosphorylation also during recovery.

**Figure 7.**
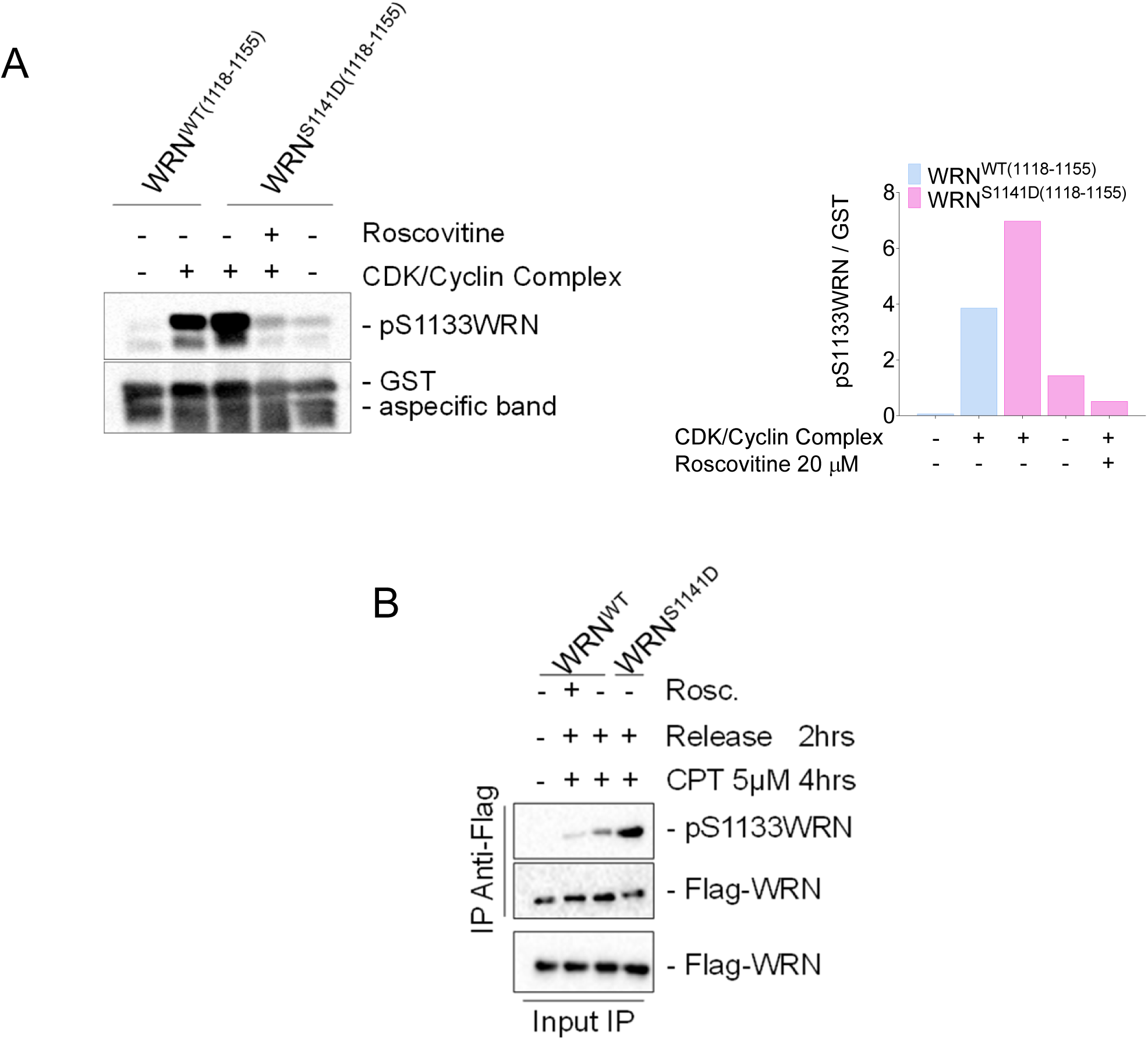
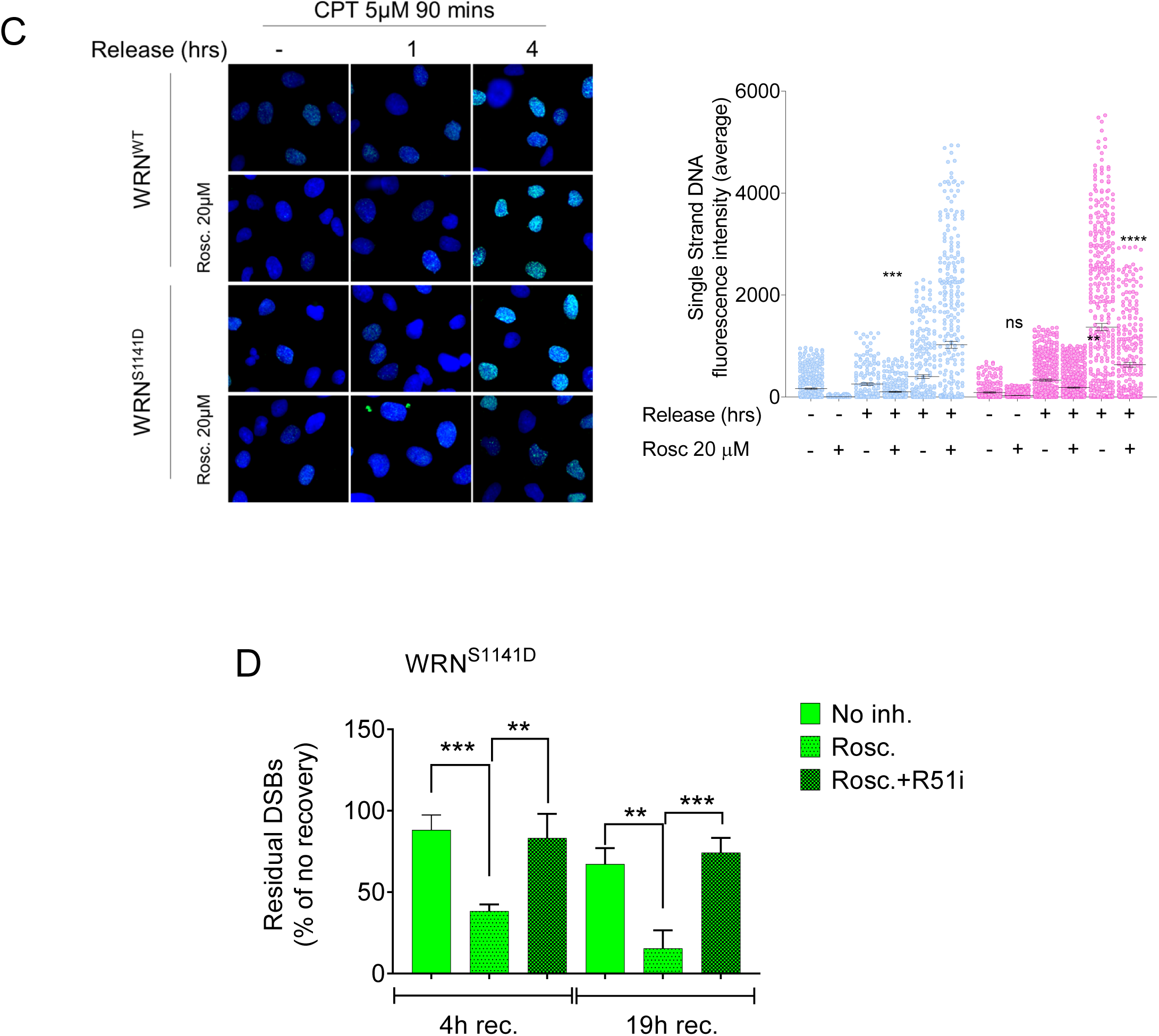
Persisting phosphorylation of WRN at S1141 leads to aberrant modification of S1131 by CDK. A) In vitro kinase assay, 2 μg of immunopurified GST-tagged WRN wild-type fragment (C-WRN^WT^) or WRN phosphomimetic mutant fragment (C-WRN^S1141D^) were phosphorylated in vitro using CDK/Cyclin complex, and the experiment was performed treated with 20μM Roscovitine. Immonoblotting was used to analyse WRN phosphorylation level in different WRN fragments using an anti-pS1133WRN antibody. The graph shows the pS1133WRN level in each experimental point. B) WRN was immunoprecipitated from cells transiently transfected with Flag-WRN wild-type (WRN^WT^) or its phosphomimetic mutant and treated 5μM CPT for 4hrs followed by release for 2 hrs treating with 20μM Roscovitine. Nine-tenth of IPs was analysed by Western Blotting (WB) with the anti-pS1133WRN antibody, while 1/10 was analysed by anti-Flag antibody. Input represents 1/50 of the lysate. Anti-Flag antibody was used to verify transfection. C) WS-derived cell lines complemented with different WRN mutants were labelled, treated with CPT and IdU/ssDNA assay was performed. The dot plot shows the mean intensity of IdU/ssDNA staining for single nuclei measured from three independent experiments (*n*=300, each biological replicate), data are presented as mean±SE. Representative images of IdU/ssDNA-stained cells are shown. Statistical analysis was performed by Anova test (\*\*\*\**P*<0.0001; *** = *P*<0.001; ns= not significant; *n*=300, each biological replicate). D) WS-derived SV40-trasformed fibroblasts were complemented with phopshomimetic WRN mutant, treated with 5 μM CPT for 1 h and allowed to recover in the presence or not of the different inhibitors as indicated. The presence of DSBs was evaluated by the neutral Comet assay. In the graph, data are presented as percent of residual DSBs from three independent experiments normalised against the value of the 0h recovery. Statistical analysis was performed by Anova test (*** = *P*<0.001; ** = *P*<0.01, *n*=300, each biological replicate).

To determine if the concomitant phosphorylation of WRN at Ser1133 and Ser1141 could be responsible for the abnormal exposure of nascent ssDNA observed at 4h of recovery in the S1141D-WRN mutant, we exposed cells to CPT and then to Roscovitine during recovery to inhibit any CDK activity (Fig. 7C). Analysis of ssDNA by the IdU/ssDNA assay revealed that, in wild-type cells, inhibition of CDKs during recovery increased the formation of ssDNA, while it largely suppressed the ssDNA accumulated in cells expressing the S1141D-WRN mutant. To verify whether inhibition of CDKs in cells expressing S1141D-WRN could also restore the repair of CPT-induced DSBs, we performed neutral Comet analysis in cells recovered in the presence or not of Roscovitine (Fig. 7D). Interestingly, inhibition of CDKs after treatment during recovery significantly suppressed the level of unrepaired DSBs detected at 4h of recovery from CPT in cells expressing the S1141D-WRN, restoring the RAD51-dependent repair.

Collectively, these results demonstrate that aberrant metabolism CPT-induced DSBs and their deficient repair depend on unscheduled phosphorylation of Ser1133 of WRN by CDK activity in the presence of the mutation mimicking constitutive ATR-dependent modification at Ser1141.

## DISCUSSION

Here, using regulation-defective mutants, we demonstrate that WRN is critical for correct execution of long-range end-resection and for post-synaptic metabolism of RAD51 nucleofilaments. WRN undergoes ordered and hierarchic phosphorylation by CDK1, ATM and ATR. Each of these events specifically regulates a defined stage during the repair of DSBs at replication forks, acting as a molecular switch. Unscheduled phosphorylation of WRN leads to abnormal accumulation of ssDNA and persistence of unproductive RAD51 foci that lead to unrepaired DNA damage.

### CDK1 and ATM cross-talk to regulate WRN function during long-range resection

End-resection of DSBs is a highly regulated process in which cells need high CDK1 activity (Symington, 2016; Trovesi et al., 2013) and, consistently, WRN is phosphorylated by CDK1 to carry out its pro-resection function (Palermo et al., 2016). However, ATM-dependent phosphorylation of several proteins is also crucial for end-resection (Symington, 2016). The WRN protein is phosphorylated by ATM on multiple residues in response to replication stress or ionizing radiation (Ammazzalorso et al., 2010; Matsuoka et al., 2007). Our data show that only two S/TQ residues, Ser1058 and Ser1141, are important for DSBs repair by recombination, but with two distinct functional roles. Phosphorylation of WRN by ATM at Ser1058 is essential for long-range end-resection and, notably, requires prior phosphorylation at Ser1133 by CDK1. Interestingly, both events implicate ATM and MRE11 nuclease activity. ATM and MRE11 are involved in initiating end-resection (Garcia et al., 2011; Kijas et al., 2015; Nimonkar et al., 2011; Williams et al., 2009), thus, our findings support the possibility that WRN phosphorylation occurs only if end-resection initiates and are consistent with the observed requirement of WRN helicase activity in the long-range phase of resection (Palermo et al., 2016; Sturzenegger et al., 2014). These findings, together with the enhanced ability of ATM to phosphorylate *in vitro* the S/TQ sites of WRN in the presence of the S1133D mutation, suggest that the primary function of Ser1133 phosphorylation is to promote subsequent ATM-dependent phosphorylation at Ser1058. Moreover, abrogation of the Ser1141 phosphorylation of WRN enhances resection at DSBs and delays recruitment of RAD51. Notably, also deregulated phosphorylation of EXO1, another nuclease involved in end-resection, is able to interfere with resection (Bolderson et al., 2010; Kijas et al., 2015), suggesting that phosphorylation by ATM/ATR is a common mechanism to regulate resection.

### Phosphorylation of WRN at Ser1141 and resolution of recombination intermediates

Loss of Ser1141 phosphorylation, however, does not confer a severe hyper-resection phenotype in response to CPT treatment. Interestingly, although expression of a WRN mimicking constitutive phosphorylation at Ser1141 induces a mild suppression of ssDNA during CPT treatment, it results in a striking increase of nascent ssDNA, which can also be observed by the CRISR/Cas9 assay, and elevated RAD51 foci late during recovery from CPT, indicating that Ser1141 phosphorylation needs to be strictly regulated. Surprisingly, this remarkable enhancement of ssDNA is driven by nucleolytic processing but depends on strand invasion as it is repressed by the RAD51 inhibitor B02, which interferes with nucleofilament formation (Huang et al., 2011).

Replication-dependent DSBs are one-ended and are normally repaired by BIR (Sakofsky and Malkova, 2017). A way replication proceeds by BIR is through migration of a D-loop (Sakofsky and Malkova, 2017). Migrating D-loop, if unproductive, leads to strand rejection and extensive resection far away the break site (Ferrari et al., 2020). In yeast, Srs2 strippase and helicase activities can resolve the RAD51 intermediates during BIR, preventing accumulation of unproductive intermediates and ssDNA behind the migrating D-loop (Elango et al., 2017). Although WRN does not have RAD51 strippase activity, it can process D-loop *in vitro* (Rossi et al., 2010) and the presence of a deregulated phosphorylation at Ser1141 could interfere with this function. A dephosphorylated WRN at Ser1141 may perform dismantling of unproductive RAD51 filaments as evidenced for the related helicases BLM and RECQ5 (Hu et al., 2007; Karow et al., 2000). However, the expression of phosphomimetic S1141D-WRN does not induce a hyper-recombination phenotype but, rather, decreases recombination efficiency, arguing against this hypothesis. Moreover, inability of S1141D-WRN to repair CPT-induced DSBs is due to the accumulation of unproductive RAD51 filaments and ssDNA blocking any other DNA repair activity. Consistent with this hypothesis, the DSB repair defect of the S1141D WRN-expressing cells can be mitigated by depletion of BRCA1 and by switching CPT-induced DSBs repair from HR to NHEJ.

Interestingly, the aberrant handling of RAD51-dependent intermediates induced by the expression of the S1141D-WRN mutant does not derive directly from the phosphomimicking mutation but rather from the concomitant pathological stimulation of Ser1133 phosphorylation by CDKs. Indeed, both the abnormal accumulation of ssDNA during recovery and the persistent DSBs are reverted if CDKs are inhibited during recovery.

### Conclusions

Our data and published observations can be summarised in the model shown in Figure 8. In response to replication-dependent DSBs and after short-range resection, WRN is phosphorylated at Ser1133 by CDK1. This phosphorylation primes that at S1058 by ATM, which is the crucial regulatory event to carry out WRN-DNA2-dependent long-range resection. After long-range resection started, WRN needs to be dephosphorylated at Ser1133, and probably at Ser1058, while it is targeted by ATR at Ser1141 to limit end-resection and allow recruitment of RAD51. Then, dephosphorylation of WRN at Ser1141 contributes to properly regulate dismantling of RAD51 foci and/or resolution of the migrating D-loop during BIR. Inability to regulate or perform this ordered phosphorylation cascade, would result in either inability to sustain end-resection or to repair DSBs by HR.

**Figure 8.**
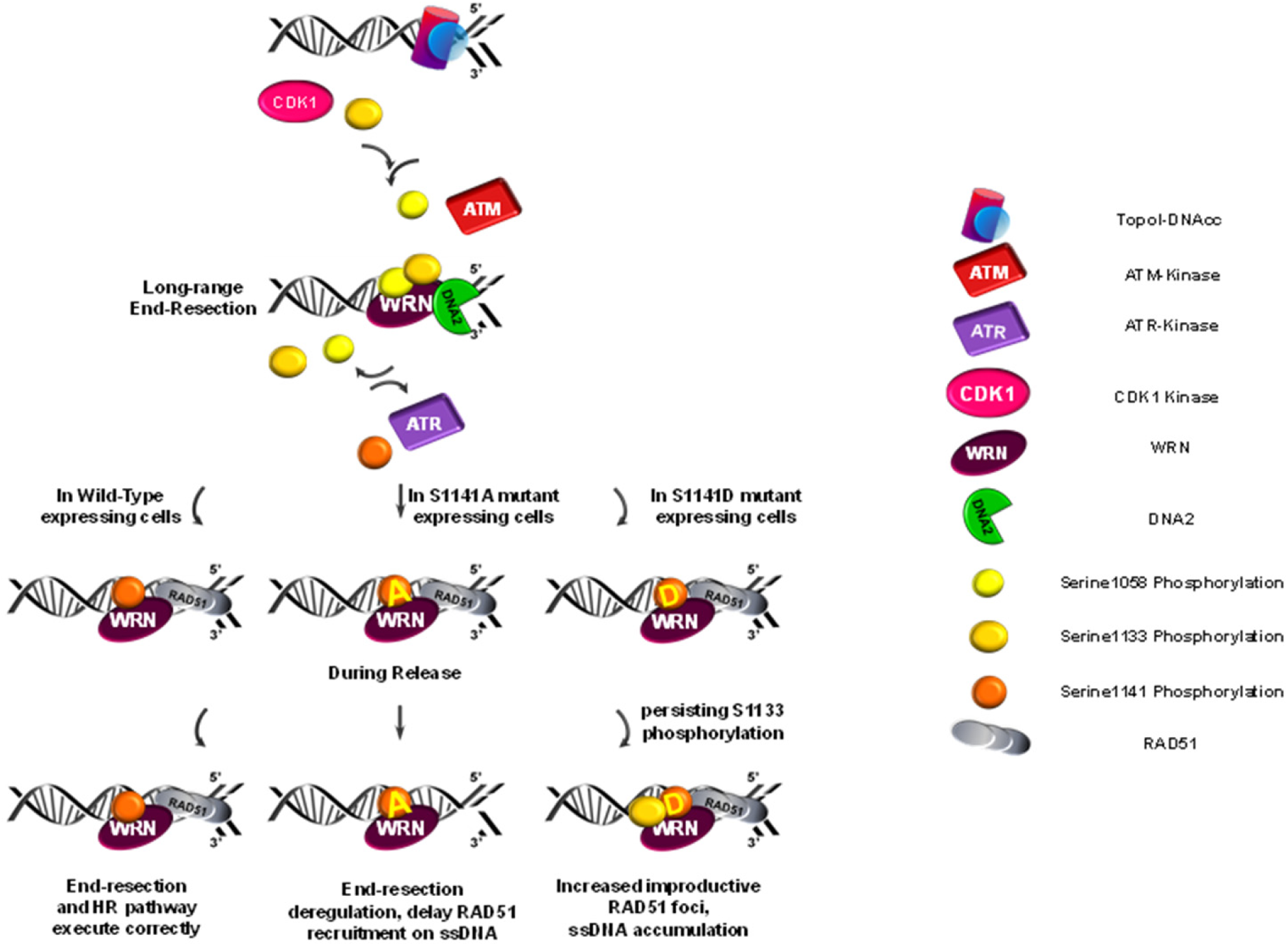
Proposed model of regulation of WRN in the repair of DSBs at the replication fork. (see text for details).

Using regulatory and separation-of-function mutants of WRN, our study reveals that the response to replication-dependent DSBs relies on multiple roles of WRN and a complex regulatory layer that controls these roles. Hence, our work indicates that WRN may act as a sort of molecular switch to put together, correctly, the end-resection and the repair stages of HR.

## MATERIALS AND METHODS

### Cell lines and culture conditions

The SV40-transformed WRN-deficient fibroblast cell line (AG11395-WS) was obtained from Coriell Cell Repositories (Camden, NJ, USA). To produce stable cell lines, AG11395 (WS) fibroblasts were transduced with lentiviruses expressing the full-length cDNA encoding wild-type WRN (WRN^WT^), S1133A-WRN (WRN^S1133A^), S1133D-WRN (WRN^S1133D^), 3AATM-WRN (WRN^3AATM^), or 3DATM-WRN (WTN^3DATM^), S1141A-WRN (WRN^S1141A^), S1141D-WRN (WRN^S1141D^). HEK293T cells were from American Type Culture Collection. HEK293TshWRN, cells were generated after transfection with pRS-puro-shWRN (5′-AGGCAGGTGTAG-GAATTGAAGGAGATCAG-3′; sequence ID: TI333414 Origene) and puromycin selection (Palermo et al., 2016). All the cell lines were maintained in Dulbecco’s modified Eagle’s medium (DMEM; Life Technologies) supplemented with 10% FBS (Boehringer Mannheim) and incubated at 37 °C in a humidified 5% CO2 atmosphere.

### Chemicals

Camptothecin (ENZO Lifesciences) was added to culture medium at 5µM if not otherwise specified to induce seDSBs. Mirin (Calbiochem), an inhibitor of MRE11 exonuclease activity, was used at 50µM. The B02 compound (Selleck), an inhibitor of RAD51 activity, was used at 27 µM.

Roscovitine (Selleck), a pan-CDKs inhibitor, was used at final concentration of 20 µM. ATM inhibitor (KU-55933, Selleck) and ATR inhibitor (VE-821, Selleck) were used at final concentration of 10 µM. An inhibitor of DNA2 nuclease (Liu et al., 2016, p. 2) was used at final concentration of 300 µM. NU7441 (Selleck), a DNAPKcs inhibitor, was used at final concentration of 1 µM, while the Ligase I/III L67 inhibitor (Sigma-Aldrich) was used at 6µM. IdU (Sigma-Aldrich) was dissolved in sterile DMEM as a stock solution 2.5 mM and stored at −20 °C.

### Oligos and plasmids

pCMV-FlagRnai-resWRN^WT^, pCMV-FlagRnai-resWRN^S1133A^ or pCMV-FlagRnai-resWRN^S1133D^ were generated ad described previously (Palermo et al., 2016). Site Directed Mutagenesis (SDM) was used to create pCMV-FlagRnai-resWRN mutant constructs. The FLAG-ATM expression plasmid was described in (Ammazzalorso et al., 2010) and the plasmid sgCAS9 was describe in (Dibitetto et al., 2018).

### Transfections

pCMV-tag2B (pFLAG) empty vector, pCMV-FlagRnai-resWRN^WT^ or other pCMV-FlagRnai-resWRN mutant constructs, as indicated, were transfected in HEK293TshWRN cell line using DreamFect (OZBioscience) following manufacturer’s instructions for 48 hours.

pCMV-tag2B (pFLAG) empty vector, pCMV-FlagRnai-resWRN^WT^, pCMV-FlagRnai-resWRN^S1058A^, pCMV-FlagRnai-resWRN^S1058D^, pCMV-FlagRnai-resWRN^S1292A^, pCMV-FlagRnai-resWRN^S1292D^ were transiently transfected in WRN-deficient fibroblast cell line using Neon transfection system 48 h prior to perform experiments. Plasmid Cas9 w/o guide were transiently transfected alone or combined with pCMV-FlagRnai-resWRN^WT^ or other pCMV-FlagRnai-resWRN mutant constructs, as indicated. BRCA1 siRNA was transfected at final concentration of 50nM using Lullaby (OZBioscience),48 hours before experiments, following manufacturer’s instructions.

### Immunoprecipitation and Western blot analysis

Immunoprecipitation experiments were performed using 2.5×10^6^ cells. RIPA buffer (0.1% SDS, 0.5% Na-deoxycholate, 1% NP40, 150mM NaCl, 1mM EDTA, 50mM Tris/Cl pH 8) supplemented with phosphatase, protease inhibitors and benzonase was used for cells lysis. One-milligram of lysate was incubated overnight at 4°C with 20 µl of Anti-Flag M2 magnetic beads (Sigma-Aldrich). After extensive washing in RIPA buffer, proteins were released in 2X electrophoresis buffer and subjected to SDS-PAGE and Western blotting.

Western blotting was performed using standard methods. Blots were incubated with primary antibodies: rabbit anti-WRN (Abcam, 1:2000); mouse anti-β-Tubulin (Sigma-Aldrich, 1:2000); rabbit anti-Lamin B1 (Abcam, 1:40000); mouse anti-GAPDH (Millipore, 1:5000), mouse anti-DDK-Flag tag (Origene, 1:2000); rabbit anti-pS/TQ (Cell Signaling Technology, 1:1000); rabbit anti-pS1141WRN (Sigma-Aldrich, 1:1000); rabbit anti-pS1133WRN (Genscript-custom, 1:10000); rabbit anti-GST (Calbiochem, 1:5000), anti-BRCA1 (Santa Cruz, 1:500). After incubation with horseradish peroxidase-linked secondary antibodies (1:30000, Jackson Immunoscience), the blots were detected using the Western blotting detection kit WesternBright ECL (Advansta) according to the manufacturer’s instructions. Quantification was performed on scanned images of blots using Image Lab software, and values shown on the graphs represent normalization of the protein content evaluated through LaminB1, or β-Tubulin-immunoblotting.

### Immunofluorescence assay

Cells were grown on 22×22 coverslip or 8 well Nunc chamber slides. To detect RAD51 foci, we performed pre-extraction for 5 min on ice in 0,5% Triton X-100 and fixed with 3% paraformaldehyde (PFA)/2% sucrose at room temperature (RT) for 10 min. After blocking in 3% bovine serum albumin (BSA) for 15 min, staining was performed with rabbit polyclonal anti-RAD51 (Abcam, 1:1000) diluted in 1% BSA/ 0,1% saponin in PBS solution, for 1 h a 37°C in a humidifier chamber. After extensive washing with PBS, specie-specific fluorescein-conjugated secondary antibody (Alexa Fluor 488-conjugated Goat Anti-Mouse IgG (H+L), highly cross-adsorbed-Life Technologies) was applied for 1h a RT followed by counterstained with 0,5µg/ml 4,6-diamidino-2-phenylindole (DAPI). Secondary antibodies were used at 1:200 dilution. Slides were analysed with Ecliplse 80i Nikon Fluorescence Microscope, equipped with a Video Confocal (ViCo) system at 60X.

### Detection of nascent single-stranded DNA by native IdU assay

To detect nascent single-stranded DNA (ssDNA), cells were grown on 22×22 coverslips in 35mm dishes. After 24h, the cells were labelled for 15 min before the treatment with 50µM IdU (Sigma-Aldrich), then treated with CPT 5µM for different time points.

For immunofluorescence cells were washed with PBS, permeabilized with 0.5% Triton X-100 for 10 min at 4°C and fixed wit 2% sucrose, 3%PFA. Fixed cells were then incubated with primary mouse anti-IdU antibody (Becton Dickinson) for 1h at 37°C in 1%BSA/PBS, followed by species-specific fluorescein-conjugated secondary antibody (Alexa Fluor 488-conjugated Goat Anti-Mouse IgG (H+L), highly cross-adsorbed-Life Technologies).

Nuclei were counterstained with 0,5µg/ml 4,6-diamidino-2-phenylindole (DAPI). Slides were analysed with Eclipse 80i Nikon Fluorescence Microscope, equipped with a Virtual Confocal (ViCo) system. For each time point, at least 100 nuclei were scored at 40×. Parallel samples either incubated with the appropriate normal serum or only with the secondary antibody confirmed that the observed fluorescence pattern was not attributable to artefacts. Quantification was carried out using ImageJ software.

### Generation of the GST-WRN fragment

DNA sequence corresponding to aa 940–1432 (C-WRN) of WRN was amplified by PCR from the pCMV-FlagWRN plasmid or from the pCMV-FlagWRN^S1133D^ mutant The PCR product were subsequently purified and sub-cloned into pGEX4T-1 vector (Stratagene) for subsequent expression in bacteria as GST-fusion proteins. The resulting vectors were subjected to sequencing to ensure that no mutations were introduced into the WRN sequence and were used for transforming BL21 cells (Stratagene). Expression of GST and GST-fusion proteins were induced upon addition of 1 mM isopropyl-D-thiogalactopyranoside (IPTG) for 2 h at 37°C. GST, and GST-C-WRN were affinity-purified using glutathione (GSH)-magnetic beads (Promega).

### In vitro Kinase assay

For kinase assay, 2 μg of immunopurified GST-tagged WRN fragment was phosphorylated in vitro by immunopurified FLAG-ATM in the presence or not of 5μM ATP for 30 minutes at 37°C. After the incubation, WRN fragments were separated from the beads and phosphorylation levels were assessed by SDS-PAGE followed by Coomassie staining and densitometric analysis by Phosphorimaging or by WB using rabbit anti-pS/TQ antibody.

### Neutral comet assay

DNA breakage induction was evaluated by Comet assay (single cell gel electrophoresis) in non –denaturing condition. Briefly, dust free frosted-end microscope slides were kept in methanol overnight to remove fatty residues. Slides were then dipped into molten low melting point (LMP) agarose al 0.5% and left to dry. Cell pellets were resuspended in PBS and kept on ice to inhibit DNA repair. Cell suspensions were rapidly mixed with LMP agarose at 0.5% kept at 37°C and an aliquot was spread onto agarose-covered surface of the slide. Agarose-embedded cells were lysed by submerging slides in lysis solution (30mM EDTA, 0.1% sodium dodecyl sulfate (SDS)) and incubated at 4°C, 1 h in the dark. After lysis, slides were washed in Tris Borate EDTA (TBE) 1X running buffer (Tris 90mM, boric acid 90 mM, EDTA 4mM) for 1 min. Electrophoresis was performed for 20 min in TBE 1X buffer at 1 V/cm. Slides were subsequently washed in distilled H_2_O and dehydrated in ice-cold methanol. Nuclei were stained with GelRed (Biotium, 1:1000) and visualized with a fluorescence microscope (Zeiss), using a 20x objective, connected to a CDD camera for image acquisition. At least 200 comets per cell line were analysed using CometAssay IV software (Perceptive instruments). To assess the amount of DNA DSB breaks, computer generated tail moment values (tail length × fraction of total DNA in the tail) were used and data from tail moments processed using Prism software. Apoptotic cells (small Comet head and extremely larger Comet tail) were excluded from the analysis to avoid artificial enhancement of the tail moment.

### In situ PLA assay for ssDNA-protein interaction

The in *situ* PLA (Merck) was performed according the manufacturer’s instruction. For nascent ssDNA-protein interaction, cells were labelled with 100µM IdU for 15 min before treatments. After treatment, cells were permeabilized with 0.5% Triton X-100 for 10 min at 4°C, fixed with 3% formaldehyde/ 2% sucrose solution for 10 min and then blocked in 3% BSA/PBS for 15 min. After washing with PBS, cells were incubated with the two relevant primary antibodies. The primary antibodies used were as follows: rabbit monoclonal anti-RAD51 (Abcam, 1:1000), and mouse monoclonal anti-BrdU/IdU (Becton Dickinson, clone b44, 1:10). We used only one primary antibody as a negative control. Samples were incubated with secondary antibodies conjugated with PLA probes MINUS and PLUS: the PLA probe anti-mouse PLUS and anti-rabbit MINUS (Merck). Incubation with all antibodies was accomplished in a humidified chamber for 1 h at 37 °C. next, the PLA probes MINUS and PLUS were ligated using two connecting oligonucleotides to produce a template for rolling cycle amplification. After amplification, the products were hybridized with red fluorescence-labelled oligonucleotide. Samples were mounted in Prolong Gold anti-fade reagent with DAPI (blue). Images were acquired randomly using Eclipse 80i Nikon Fluorescence Microscope, equipped with a Video Confocal (ViCo) system.

### Homologous recombination Reporter Assay

HEK293TshWRN cells (Palermo et al., 2016) were seeded in 6-well plates at a density of 0.5 million cells/well. The next day, pCMV-FlagRnai-resWRN^WT^ or other pCMV-FlagRnai-resWRN mutant constructs were co-transfected with the I-SceI expression vector pCBASceI and the pHPRT-DRGFP plasmid reporter using DreamFect (OZBioscience) according to the manufacturer’s instruction. pHPRT-DRGFP and pCBASceI were a gift from Maria Jasin (Addgene plasmids # 26476 and #26477).

Protein expression levels were analyzed by Western Blotting 72 h post-transfection. Cells were subjected to flow cytometry analysis at 72 h after transfection to determine the percentage of GFP-positive cells.

### Human genomic DNA extraction

The SV40-transformed WRN-deficient fibroblast cell line were grown on 6-well plates after transfection. At the indicated time points, cells where trypsinized, washed in PBS and genomic DNA was extracted by NucleoSpin^TM^ Tissue kit (Macherey-Nagel), according to the manufacturer’s instructions. The day after, 15 µl of genomic DNA (DNA concentration is around 100 ng/µl) were digested or mock with 20 units of *Bsr*GI or *Bam*HI restriction enzymes (New England BioLabs) for 5 hours at 37 °C. Digested or mock DNA was purified and 5 µl were used for the ddPCR reaction.

### Droplet Digital PCR assay

The ddPCR reaction was assembled as follows: 5 µl of genomic DNA (approximately 50 ng), 1X ddPCR^TM^ Supermix for Probes (no dUTP, Bio-Rad), 900 nM for each pair of primers, 250 nM for each probe (HEX and FAM) and dH_2_O to 20 µl per sample. We produced droplets pipetting 20 µl of the PCR reaction mix into single well of a universal DG8^TM^ cartridge for droplets generation (Bio-Rad). 70 µl of droplet generation oil were also added in each well next to the ones containing the samples. Cartridges were covered with DG8^TM^ droplet generator gaskets (Bio-Rad) and then placed into the droplet generator (QX200^TM^, Bio-Rad). After droplet generation, 40 µl of emulsion were transferred from the right well of the cartridge to a 96-well ddPCR plate (BioRad). Before PCR reaction, 96-well PCR plates were sealed with peelable foil heat seals at the PCR plate sealer machine (PX1^TM^, Bio-Rad).

PCR (T100^TM^ thermocycler, BioRad) was run using a ramp rate of 2.5°C/s between each step. First Taq polymerase was activated at 95 °C for 5 minutes and then 39 cycles of 95°C for 30 s and 58.7°C for 1 minute were made. At the end of the cycles, one additional cycle at 4 °C for 5 minutes and one at 90°C for 5 minutes were made, then temperature was held at 12°C. After the PCR, FAM and HEX fluorescence was read at the droplet reader (QX200^TM^, BioRad) using QuantaSoft^TM^ software (BioRad). For each sample the number of droplets generated were on an average of 15 000. The number of copies/µl of each target locus was determined setting an empirical baseline threshold identical in all the samples.

Calculation of Cas9 cleavage efficiency and measurement of ssDNA was performed as described in Dibitetto et al., (Dibitetto et al., 2018).

### Statistical analysis

All the data are presented as means of at least two pooled independent experiments. Statistical comparisons of WS or WRN-mutant cells to their relevant control were performed by one-sided analysis of variance (ANOVA test: Comet assays), or Mann-Whitney test (ssDNA, PLA and other experiments) using the built-in tools in Prism 8 (GraphPad Inc.). P < 0.05 was considered as significant. Statistical significance was always denoted as follow: ns = not significant; * p < 0.05; ** p < 0.01; *** p < 0.001; **** p < 0.0001. Any specific statistical analysis is reported in the relevant legend.

## ACKNOWLEDGMENTS

This work was supported by Associazione Italiana per la Ricerca sul Cancro (AIRC) to PP (IG n. 21428) and to A.F. (IG n. 19971), and to A.P. (IG. 19917) and by Telethon grant n. 12144 to PP.

## AUTHOR CONTRIBUTIONS

V.P. performed experiments to analyse WRN phosphorylation, end-resection, protein relocalisation and analysis of DNA damage. P.P. performed *in vitro* ATM kinase assays. M.S. performed flow cytometry analysis. E.M. performed the analysis of DSBs extinction with inhibitors of repair pathway and contributed to the analysis of WRN phosphorylation. M.S. contributed to analysis of WRN phosphorylation and to the cell biology experiments. B.P. performed analysis of end-resection with inhibitors of nucleases. P.V. participated to cell cycle analysis. We thank Alda Corrado for helping to perfom the molecular analysis of end-resection. V.P., F.M., A.P. and M.S. analysed data and contributed to designing the experiments and writing the paper. A.F., F.M, A.P. and P.P designed experiments, analysed data and wrote the paper. All authors approved the paper.

