## Supplementary Figures and Legends for "Switch-Like Phosphorylation of WRN Integrates End-Resection with RAD51 Metabolism at Collapsed Replication Forks"

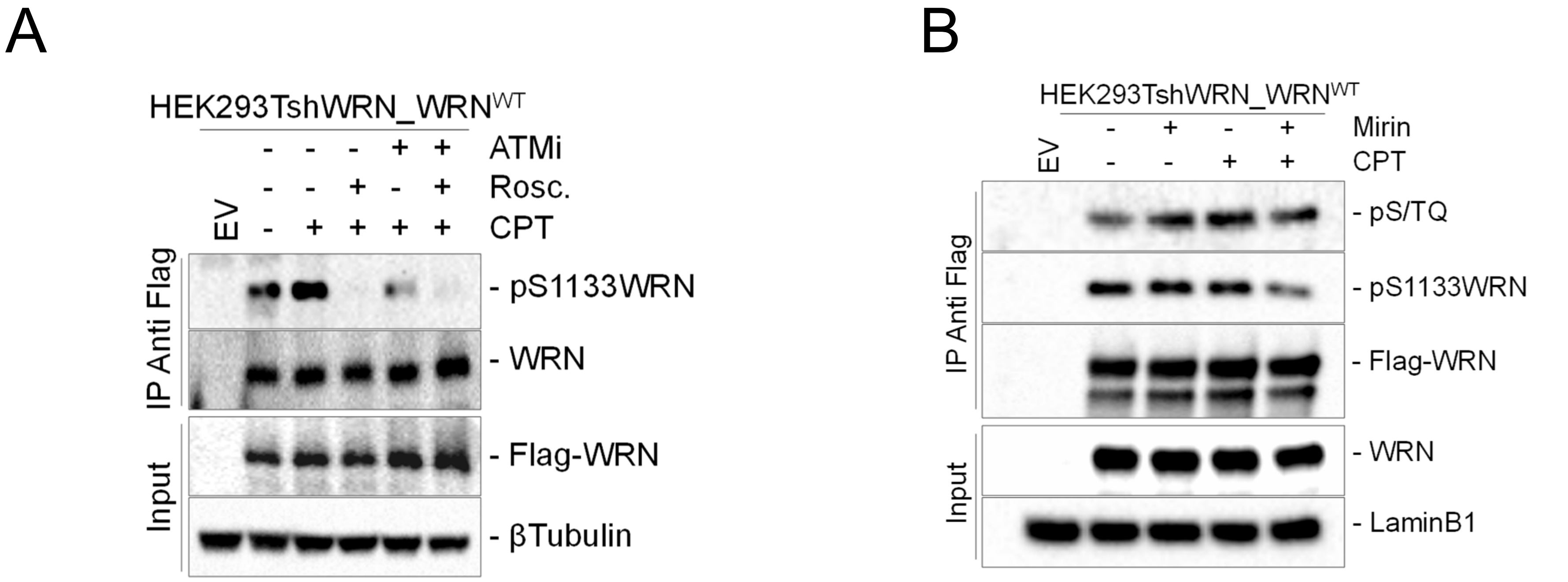


**Figure S1. Phosphorylation of WRN requires MRE11 nuclease activity**

A) Cells were treated with 10μM ATMi or 20μM Roscovitine alone or in combination, and with 5μM CPT for 4 hours. Cells were lysed and WRN protein was immunoprecipitated with anti-Flag-conjugated beads. Nine-tenth of IPs were analysed by WB with the anti-pS1133WRN antibody, while 1/10 was detected by anti-Flag antibody, as indicated. One-fiftieth of the lysate (input) was blotted with an anti-Flag antibody to verify transfection. An anti-LaminB1 antibody was used as loading control. B) Cells were treated with 50μM Mirin and 5μM CPT for 4 hours. Cells were lysed and WRN protein was immunoprecipitated with anti-Flag-conjugated beads. Nine-tenth of IPs were analysed by WB with the anti-pS1133WRN antibody and the anti-pS/TQ antibody, while 1/10 was detected by anti-Flag antibody, as indicated. One-fiftieth of the lysate (input) was blotted as in “A”.

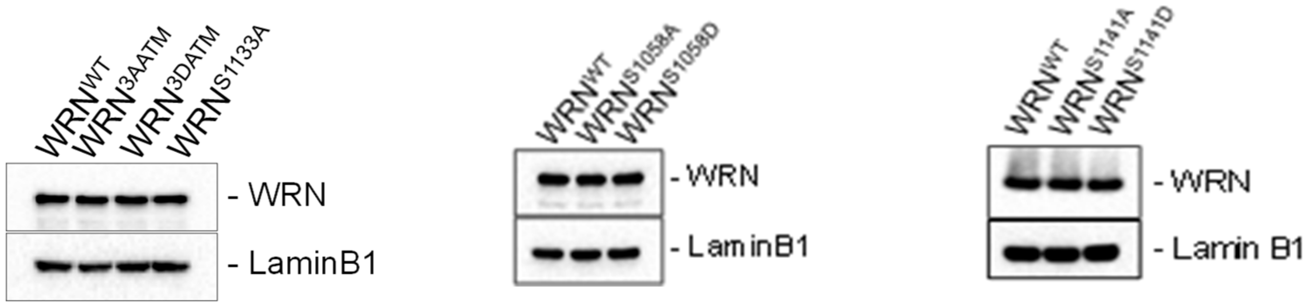


**Figure S2. Analysis of WRN level**
WS-derived SV40-trasformed fibroblasts stably expressing the wild-type form of WRN or the indicated mutant form of WRN were analysed for the protein level by WB using anti-WRN antibody. An anti-LAMINB1 was used as a loading control.

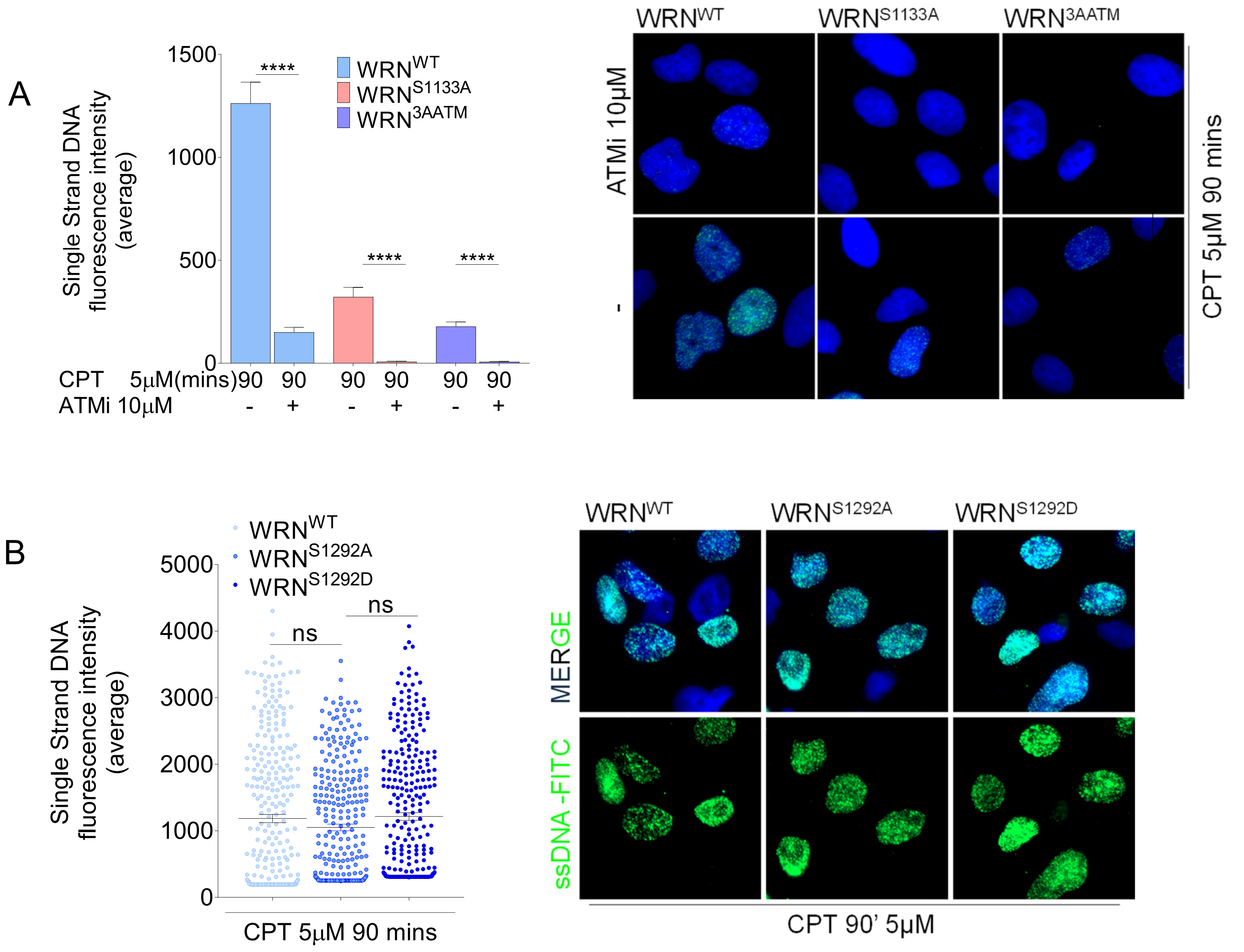


**Figure S3. Analysis of ssDNA formation**

A) WS-derived SV40-trasformed fibroblasts stably expressing the wild-type form of WRN, the 3AATM, or S1133A mutant were treated with CPT in combination or not with 10μM ATMi as indicated. The presence of ssDNA was analysed by non-denaturing IdU/ssDNA assay. The graph shows the mean intensity of IdU/ssDNA staining for single nuclei measured from three independent experiments (n=300, each biological replicate), data are presented as mean ± SE. Representative images of IdU/ssDNA-stained from CPT-treated cells are shown. DAPI was used to counterstain nuclei. Statistical analysis was performed by the ANOVA test (**** = p<0.0001). B) WS-derived SV40-trasformed fibroblasts transiently expressing WRN mutants, as indicated, were labelled, treated and IdU/ssDNA assay was performed. The dot plot shows the mean intensity of ssDNA staining for single nuclei measured from two independent experiments (n=300, each biological replicate), data are presented as mean ± SE. Representative images of IdU/ssDNA-stained from CPT-treated cells are shown. Statistical analysis was performed by the ANOVA test (**** = P<0.0001; ** = P<0.01; ns = not significant).

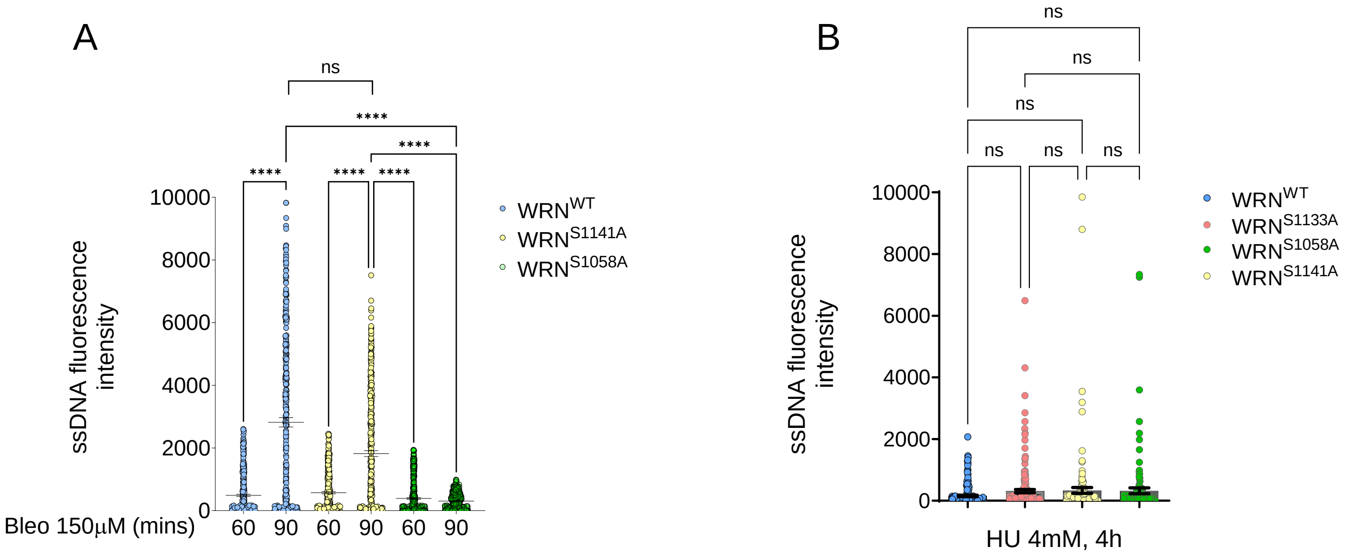


**Figure S4. Analysis of ssDNA formation**

A) WS-derived SV40-trasformed fibroblasts stably expressing the wild-type form of WRN, the S1058A, or S1141A mutant were treated with Bleomycin (Bleo) as indicated. The presence of ssDNA was analysed by non-denaturing IdU/ssDNA assay. The graph shows the mean intensity of IdU/ssDNA staining for single nuclei measured from three independent experiments (at least n=300, minum 100 each biological replicate), bars represent mean ± SE. Statistical analysis was performed by the ANOVA test (**** = p<0.0001). B) WS-derived SV40-trasformed fibroblasts transiently expressing WRN mutants, as indicated, were labelled, treated with HU as indicated and IdU/ssDNA assay was performed. The dot plot shows the mean intensity of ssDNA staining for single nuclei measured from two independent experiments (n=300, 100 each biological replicate), bars are mean ± SE. Statistical analysis was performed by the ANOVA test (ns = not significant).

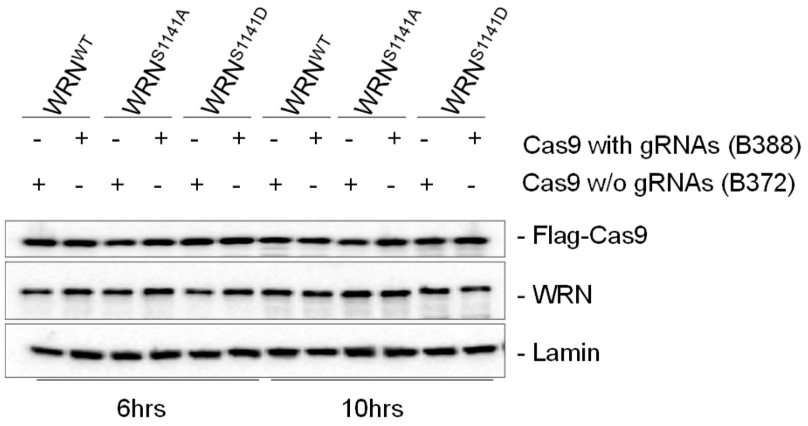


**Figure S5. Analysis of WRN and Cas9 expression levels with or without guides**
WS-derived SV40-trasformed fibroblasts stably expressing the wild-type form of WRN or the indicated mutant form of WRN were transiently transfected with the plasmid expressing Cas9 with or without the indicated sgRNAs. The expression level of Cas9 was analysed by WB using anti-Flag antibody. An anti-WRN antibody was used to analyse WRN expression and anti-LAMINB1 was used as a loading control.

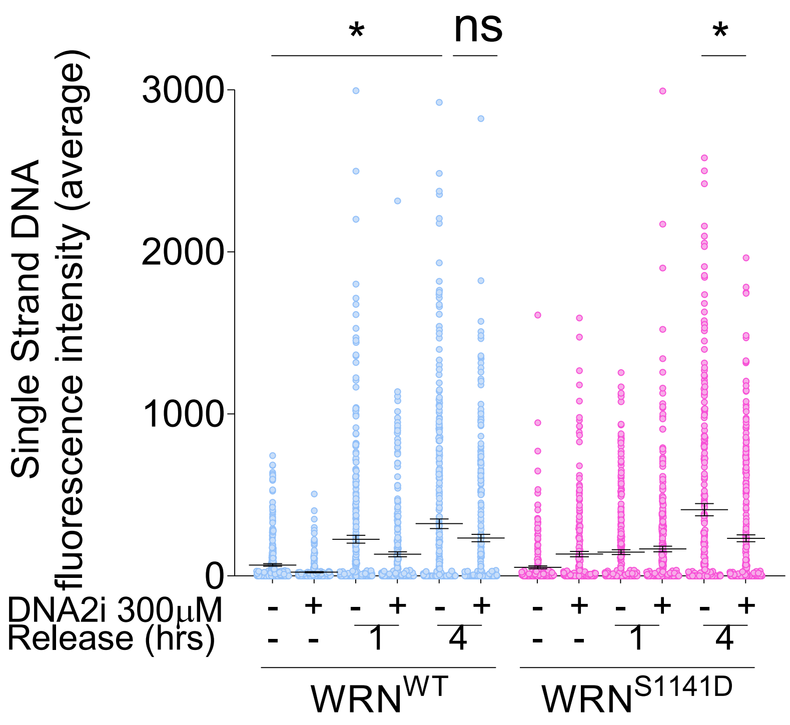


**Figure S6. Kinetics of ssDNA formation in cells treated with DNA2i**

WS cell line complemented with WRN wild-type (WRNWT) or S1141D-WRN were treated with CPT and recovered at different time points, as indicated. The graph shows the mean intensity of IdU/ssDNA staining for single nuclei measured from three independent experiments (n=300, each biological replicate), data are presented as mean ± SE. Statistical analysis was performed by the ANOVA test (* = P<0.05; ns = not significant).

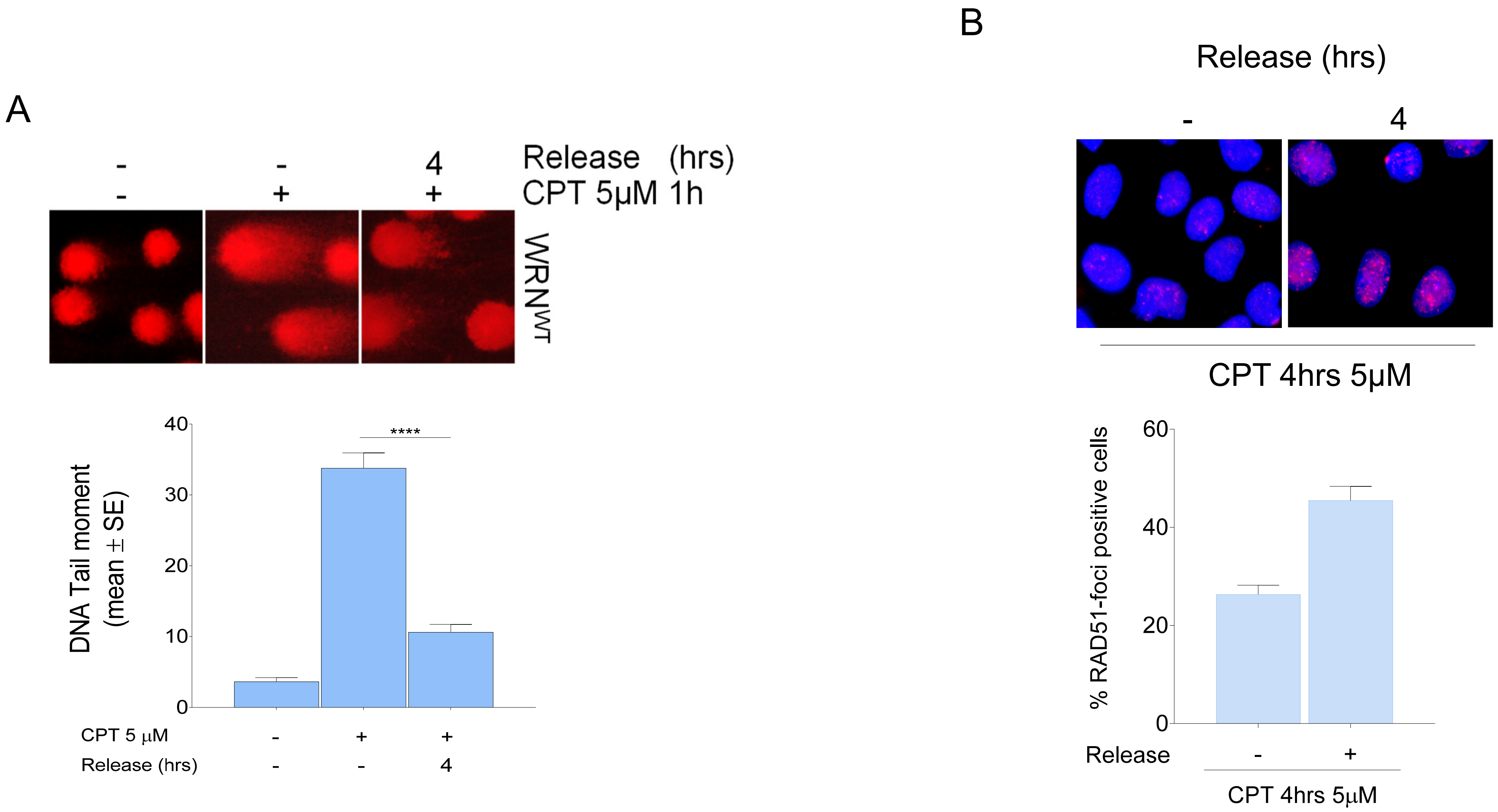


**Figure S7. Analysis of DNA repair and RAD51 recruitment during recovery from CPT**
A) WS-derived SV40-trasformed fibroblasts stably expressing the wild-type form of WRN, were treated as indicated and DSBs evaluated by neutral Comet assay. The graph shows the percentage mean tail moment as obtained from two independent experiments (n=200, each biological replicate), data are presented as mean ± SE. Representative images are shown. Statistical analysis was performed by the ANOVA test (**** = P<0.0001).
B) Cells were treated as in "A" and analysed for RAD51-foci staining by IF. The graph shows the percentage of RAD51-foci positive cells as obtained from two independent experiments (n=200, each biological replicate), data are presented as mean ± SE. Representative images are shown.

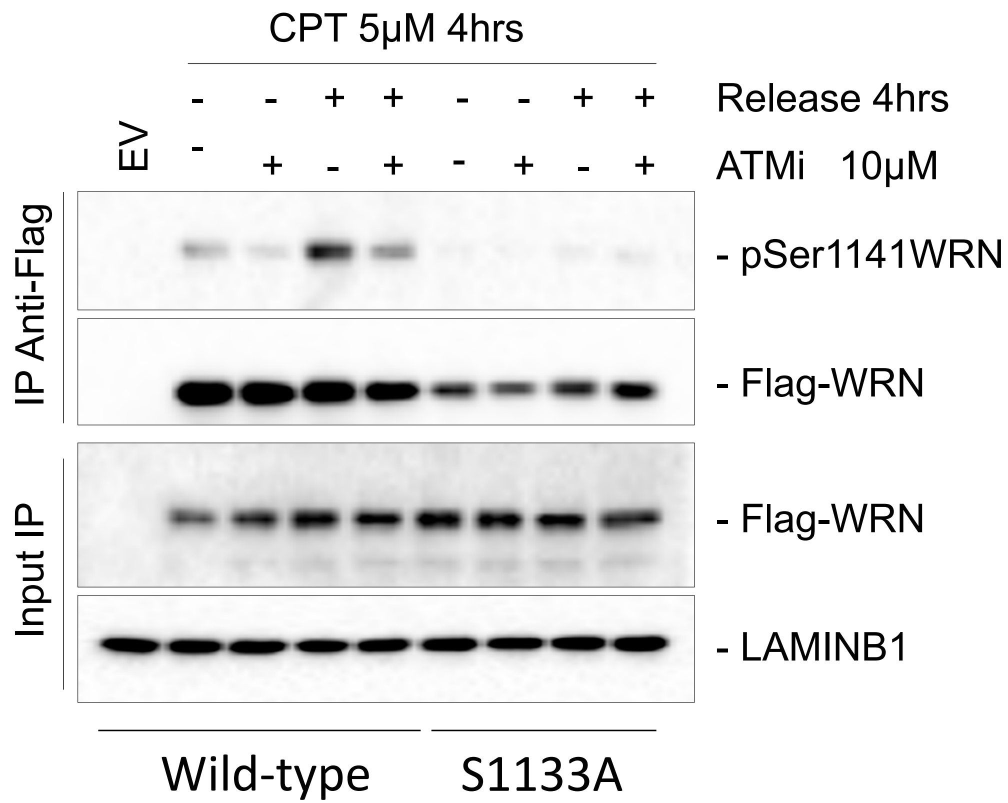


**Figure S8. Phosphorylation of WRN at S1141 requires prior phosphorylation at S1133**
Cells were treated with 10µM ATMi and with 5µM CPT for 4 hours and then released for 4h. Cells were lysed and WRN protein was immunoprecipitated with anti-Flag-conjugated beads. Nine-tenth of IPs were analysed by WB with the anti-pS1141WRN antibody, while 1/10 was detected by anti-Flag antibody, as indicated. One-fiftieth of the lysate (input) was blotted with an anti-Flag antibody to verify transfection. An anti-LaminB1 antibody was used as loading control.

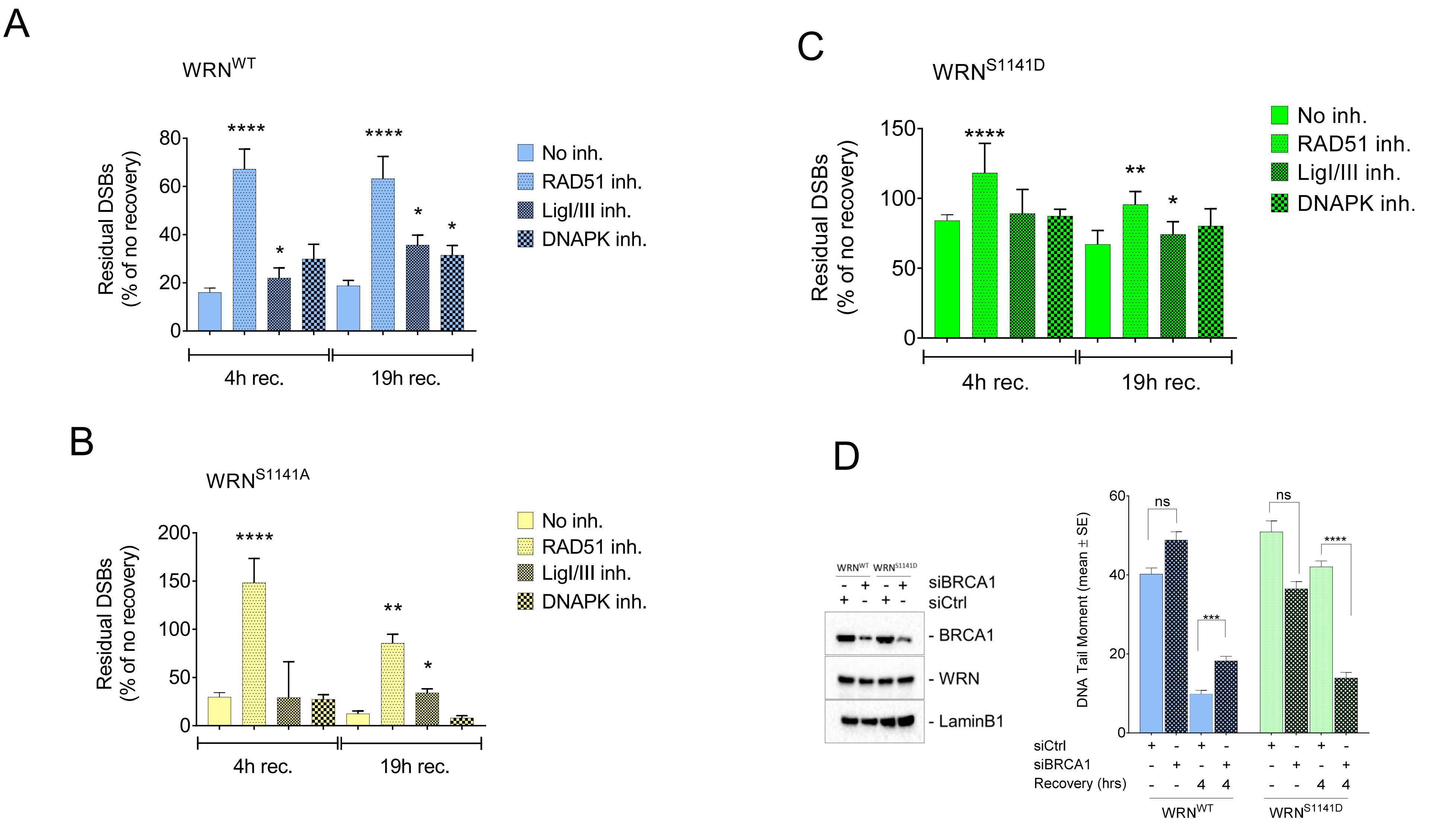


**Figure S9. Analysis of DNA repair during recovery from CPT**

A) WS-derived SV40-trasformed fibroblasts stably expressing the wild-type form of WRN or the two S1141 WRN mutants (B and C), were treated with 5μM CPT for 4 hours and DSBs evaluated by neutral Comet assay during recovery in the indicated DNA repair inhibitor, as indicated. The graph shows the percentage mean tail moment as obtained from two independent experiments (n=200, each biological replicate), data are presented as mean ± SE. D) After transfection with siCtrl or siBRCA1 oligos, cells expressing WRN wild-type or S1141D mutant were treated as in "A" and analysed for the repair of DSBs by neutral Comet assay at 4h of recovery from CPT. WB shows the depletion level normalised against the loading control LaminB1. The graph shows the percentage mean tail moment as obtained from two independent experiments (n=200, each biological replicate), data are presented as mean ± SE. Statistical analysis was performed by the ANOVA test (**** = P<0.0001; *** = P<0.001; ** = P<0.01).
